# *Neuron Navigator 1* Regulates Learning, Memory, and the Response to Multiple Potentially Addictive Drugs

**DOI:** 10.1101/2022.11.21.517383

**Authors:** Jared R. Bagley, Yalun Tan, Wan Zhu, Zhuanfen Cheng, Saori Takeda, Zhouqing Fang, Ahmed Arslan, Meiyue Wang, Yuan Guan, Lihua Jiang, Ruiqi Jian, Feng Gu, Isabel Parada, David Prince, J. David Jentsch, Gary Peltz

## Abstract

Genetic variation accounts for much of the risk for developing a substance use disorder (SUD). Inbred mouse strains exhibit substantial and heritable differences in the extent of voluntary cocaine intravenous self-administration (IVSA). Computational genetic analysis of IVSA data obtained from an inbred strain panel identified *Nav1,* a member of the neuron navigator family that regulates dendrite formation and axonal guidance, as a candidate gene. To test this hypothesis, we generated and characterized *Nav1* knockout (KO) mice. *Nav1* KO mice exhibited increased cocaine intake during IVSA testing. Surprisingly, *Nav1* KO mice also displayed a reduced susceptibility to become opioid dependent or develop opioid-induced hyperalgesia after chronic morphine administration, and had impaired spatial learning/memory. Immunohistochemistry and electrophysiology studies revealed that inhibitory synapse density in the cortex of *Nav1* KO mice was reduced, and excitatory synaptic transmission was increased in the *Nav1* KO cortex and hippocampus. Transcriptomic analysis revealed that *Nav1* KO mice had a marked increase in excitatory neurons in a deep cortical layer. Collectively, our results indicate that *Nav1* regulates learning, memory, and the response to multiple addictive drugs, and that changes in the excitatory and inhibitory synaptic balance in the cortex and hippocampus could possibly mediate these phenotypic effects.

## Introduction

While a substantial proportion of the risk for developing a substance use disorder is genetic (*1, 2*), the genes and alleles that influence susceptibility to drug addiction remain largely unknown. Just as in the human population, inbred mouse strains exhibit substantial and heritable differences in their responses to commonly used drugs with high abuse potential, which has enabled murine models for many of the behaviors observed in those with a SUD to be generated (*3*). Inbred mouse strains have been analyzed to identify genetic factors affecting addiction-related behaviors and responses to multiple types of drugs with high abuse potential (*4-8*). While we do not expect that murine studies will be likely to identify the same genetic factors that are operative in humans, the neurobiological mechanisms that underlie the risk for a SUD are likely to converge in mice and humans (*3*). Hence, characterizing genetic factors that affect the responses of inbred mouse strains to these drugs could (i) increase our understanding of the neurobiological pathways that are impacted by them; (ii) reveal how addiction-related behaviors are generated; and (iii) could help to generate new approaches for preventing drug addiction in humans (*9*). Among the various rodent assays, the cocaine intravenous self-administration (**IVSA**) assay is considered a gold-standard assay for studying cocaine-related aspects of behavior, neurobiology, and genetics in rodent populations (*10-12*). Mice are fitted with a jugular catheter and placed in an operant conditioning box where they actuate a lever to trigger cocaine infusions, which enables the extent of cocaine IVSA to be measured. The rate of cocaine IVSA reflects the reinforcing potential of cocaine (*13*), and inter-strain differences in the magnitude of cocaine IVSA reflect the propensity of a strain to misuse cocaine (*14-16*).

We analyzed a murine genetic model for cocaine IVSA and identified a candidate gene (*Nav1*) that could regulate cocaine responses. Analysis of a *Nav1* KO mouse confirmed that Nav1 had a strong effect on the level of voluntary cocaine consumption; and that it also impacted other a strong effect on the level of voluntary cocaine consumption; and that it also impacted other addiction-related traits that include opioid responses, food reinforcement, anxiety, and learning and memory. Some potential insight into the mechanism for its multiple effects was provided by finding that the *Nav1* KO altered the excitatory/inhibitory synaptic balance in hippocampal and cortical brain regions.

## Results

### Identification of Nav1 as a candidate gene affecting cocaine IVSA

Voluntary cocaine IVSA was examined over a 10 day period (0.5 mg/kg/infusion) in adult male mice of 21 inbred mouse strains. During the last 3 days of testing, the strains exhibited very different levels of cocaine IVSA (range 0 to 65 infusions), and the heritability for the inter-strain differences in this measurement was 0.72 (**Fig. 1A**). To identify genetic factors, the cocaine IVSA data was analyzed by haplotype-based computational genetic mapping (HBCGM) (*17-19*), and the allelic patterns within multiple genes that were co-localized within a region on chromosome 1 (134-137 MB) were most strongly associated with the inter-strain differences in cocaine IVSA (Fig. 1).

**Figure 1.**
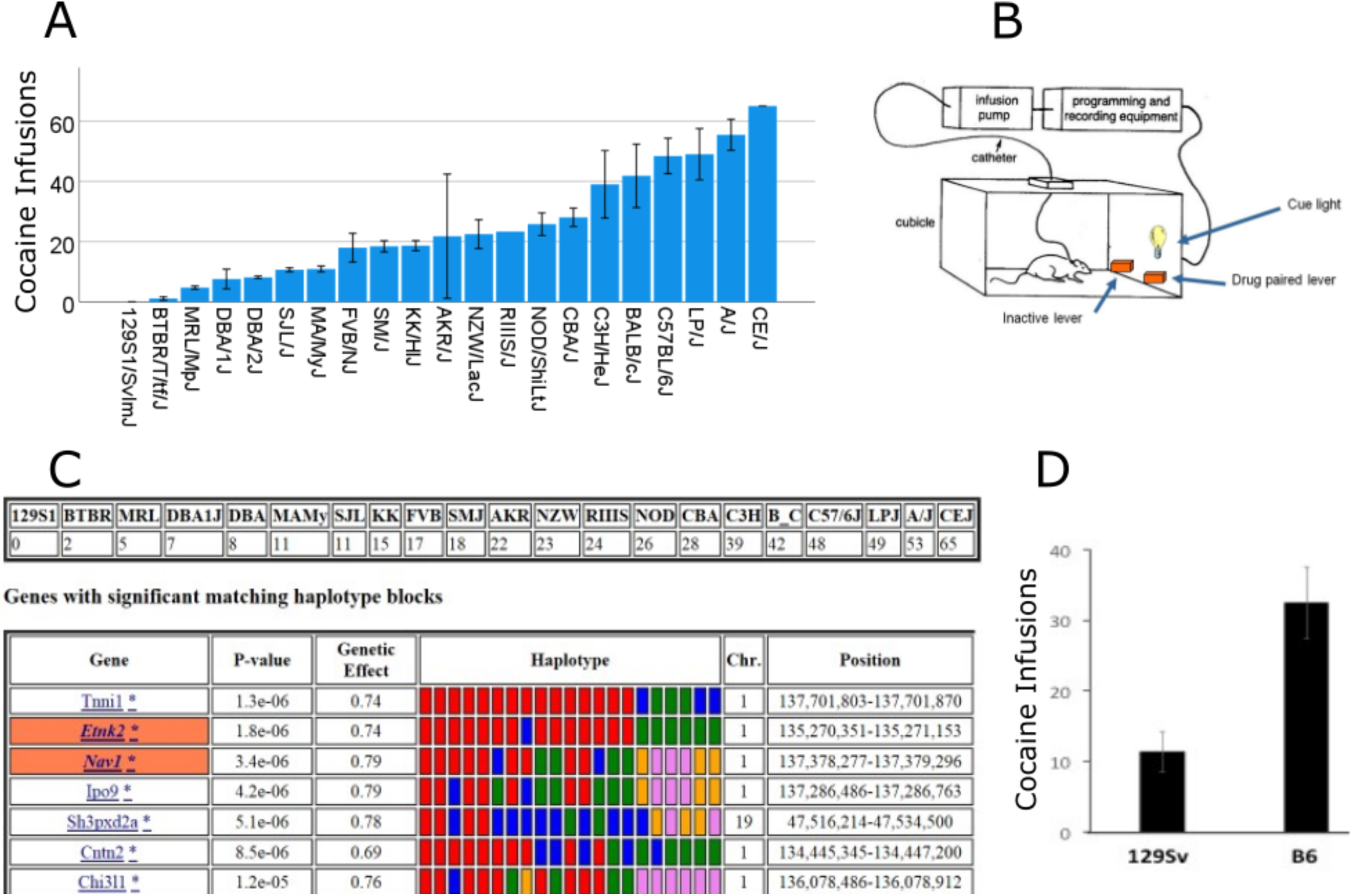
**(A)** The average number of cocaine infusions earned during the last 3 days of a 10-day cocaine IVSA session was measured for each of 21 inbred strains. Each bar shows the strain mean ± SEM (n=1-6 mice per strain). **(B)** A diagram of the cocaine IVSA assay. **(C)** The top panel shows the average number of cocaine infusions for each strain, and the bottom panel shows the top 7 genes identified by HBCGM analysis of this data. The gene symbol, genetic effect size, chromosomal position, and the p values for the genetic association are shown. If the box surrounding the gene symbol has an orange color, it indicates that the haplotype block has a SNP allele causes a significant change in the protein sequence. In the haplotype box, the haplotypic pattern is shown as colored rectangles that are arranged in the same order as the input data shown above; strains with the same-colored rectangle have the same haplotype. **(D)** The mean ± SEM for the number of cocaine infusions (y-axis) self-administered by strains with different *D198E* cSNP alleles in *Nav1* were substantially different (p=0.007). The 13 strains (CE/J, LP/J, Balbc/J, C3H/HeJ, CBA/J/ NOD/J, RIIIS, SMJ, FVB, Mamy, BTBR) with the C57BL/6 (*D198)* allele self-administered an average of 32.5 ±5.2 cocaine infusions while the 8 strains (NZW, AKR, KK, SJL, DBA2, DBA1, MRL, 129Sv1) with the 129Sv1 (*E198)* allele self administered an average of 11.3 ±2.8 cocaine infusions.

Only two of these genes (*Nav1, Etnk2*) had SNP alleles that altered its predicted amino acid sequence. However, *Nav1* was the only one of these genes with a high level of mRNA expression in key brain regions, and its allelic pattern was not associated with population structure (*20*) (**Table S1**). There were 2163 SNPs within or near (± 10 kB) the *Nav1* gene, which included 20 synonymous SNPs, and 3 cSNPs (*D198E, P1366L, A911V)* that altered the Nav1 amino acid sequence (**Table S2**). Also, two SNPs (*D198E, A911V)* were within predicted sumoylation (193-202 -EAAVS**D**DGKS) and phosphorylation (910-917 -T**A**PSEEDT) motifs. While we certainly do not know the specific *Nav1* alleles that contribute to the different levels of cocaine IVSA exhibited by the inbred strains, it is noteworthy that the level of cocaine IVSA was correlated with the *Nav1 D198E* allele of an inbred strain (p=0.007) (Fig. 1D).

Several other factors suggested that genetic variation within *Nav1* could affect IVSA. (i) *Nav1* belongs to the neuron navigator gene family, and it is expressed predominantly in the nervous system (*21*). Nav1 protein plays a role in neuronal development and in the directional migration of neurons (*22*). It regulates neurite outgrowth through effects on cytoskeletal remodeling (*23*), which are processes that have been associated with addiction-induced changes in brain (*24*). (ii) Multiple *Nav1* mRNA isoforms are produced by alternative splicing (*21*), which provides another mechanism by which allelic differences could impact a phenotype. (iii) A prior HBCGM analysis led to the discovery that alleles within another axonal guidance protein (*Netrin 1*) affected multiple maladaptive responses to opiates (*7*). (iv) Analysis of striatal *Nav1* mRNA expression levels in a BXD recombinant inbred strain panel indicated that *cis* acting alleles regulate its expression (**Fig. S1**). (v) When the *Nav1* mRNA expression QTL was evaluated for correlation with the entire database of phenotypes measured in this panel of recombinant inbred strains, the highest correlation was an inverse one with striatal *dopamine receptor D2 (Drd2)* mRNA levels. This correlation is specific in that striatal *Drd1* mRNA expression is positively associated with *Nav1* mRNA levels (Fig. S1) and is noteworthy since low Drd2 expression and function was repeatedly identified as a biomarker for addiction vulnerability in animals and humans (*25-29*). All of these features suggest that *Nav1* allelic variation could impact Drd2 dopamine-dependent signaling within the striatum, and thus, responses to potentially addictive drugs.

### Generation and characterization of a Nav1 knockout (KO) mouse

Due to its large size, the presence of many SNPs within *Nav1*, and the multiple mechanisms by which *Nav1* alleles could impact a genetic trait, the genetic hypothesis was tested by producing a homozygous *Nav1* KO mouse on a strain background (C57BL/6J) that exhibited a moderate level of cocaine IVSA (*30*) (**Fig. S2**). Single molecule FISH analysis indicated that full length *Nav1* mRNA is expressed in C57BL/6J (but not in *Nav1* KO) brain tissue, while the *Nav1* KO transcript is exclusively expressed in *Nav1* KO mice (**Fig. S3**). Proteomic analysis confirmed that Nav1 protein was absent in brain tissue obtained from *Nav1* KO mice (see methods). MRI brain scans revealed that *Nav1* KO mice did not have gross neuroanatomical abnormalities. *Nav1* KO mice had a slightly smaller overall brain volume, which was consistent with their smaller body size (vs. C57BL/6J mice). Similarly, after normalization of hippocampal volume to overall brain size, there was no significant difference in their relative hippocampal volume (vs. C57BL/6J mice) (**Fig. S4**).

### The Nav1 KO effects cocaine IVSA

A between-subjects dose-response design was used to compare the level of cocaine IVSA for wildtype, heterozygous *Nav1* (**HET**) and homozygous *Nav1* KO mice, and the data were analyzed by mixed ANOVA. The results for the effect that an evaluated variable (i.e., mouse genotype, sex, or the effect of an individual session) had on the number of cocaine infusions are reported as an F-statistic [F(variation among sample means / variation within sample)] and a p-value. The variables that had a signficant effect (or where interactions between variables were found) were further evaluated by performing pairwise comparisons of that variable (i.e., genotype or sex) with the cocaine IVSA results obtained at each session, and results are reported as a p-value for the effect of that variable on the number of cocaine infusions. Analysis of this data indicated that the genotype [F[2,113)=8.5, p<0.001] and sex [F1,113)=7.2, p=0.008] of the mouse had a clear effect on the number of cocaine infusions earned. A session-genotype interaction [F[8.6,483.3)=2.8, p=0.003], along with a main effect of session [F4.3, 483.3)=4.4, p=0.001] was also found. The pairwise comparisons of the cocaine IVSA results obtained at each session for mice of each genotype indicated that *Nav1* KO mice earned more infusions than HET *Nav1* mice (p<0.05) following session 1 and more infusions than wildtype mice following session 2 (except for session 7 (p=0.052)). These results indicate that *Nav1* KO mice self-administered more cocaine infusions, their increased consumption was observed across all cocaine doses tested, their increased level of consumption appeared during the early period of cocaine IVSA acquisition and was maintained throughout the 10 days of testing (**Fig. 2** **A, B**). The sex effect indicated that males consumed more cocaine than females across all doses and genotypes tested.

**Figure 2.**
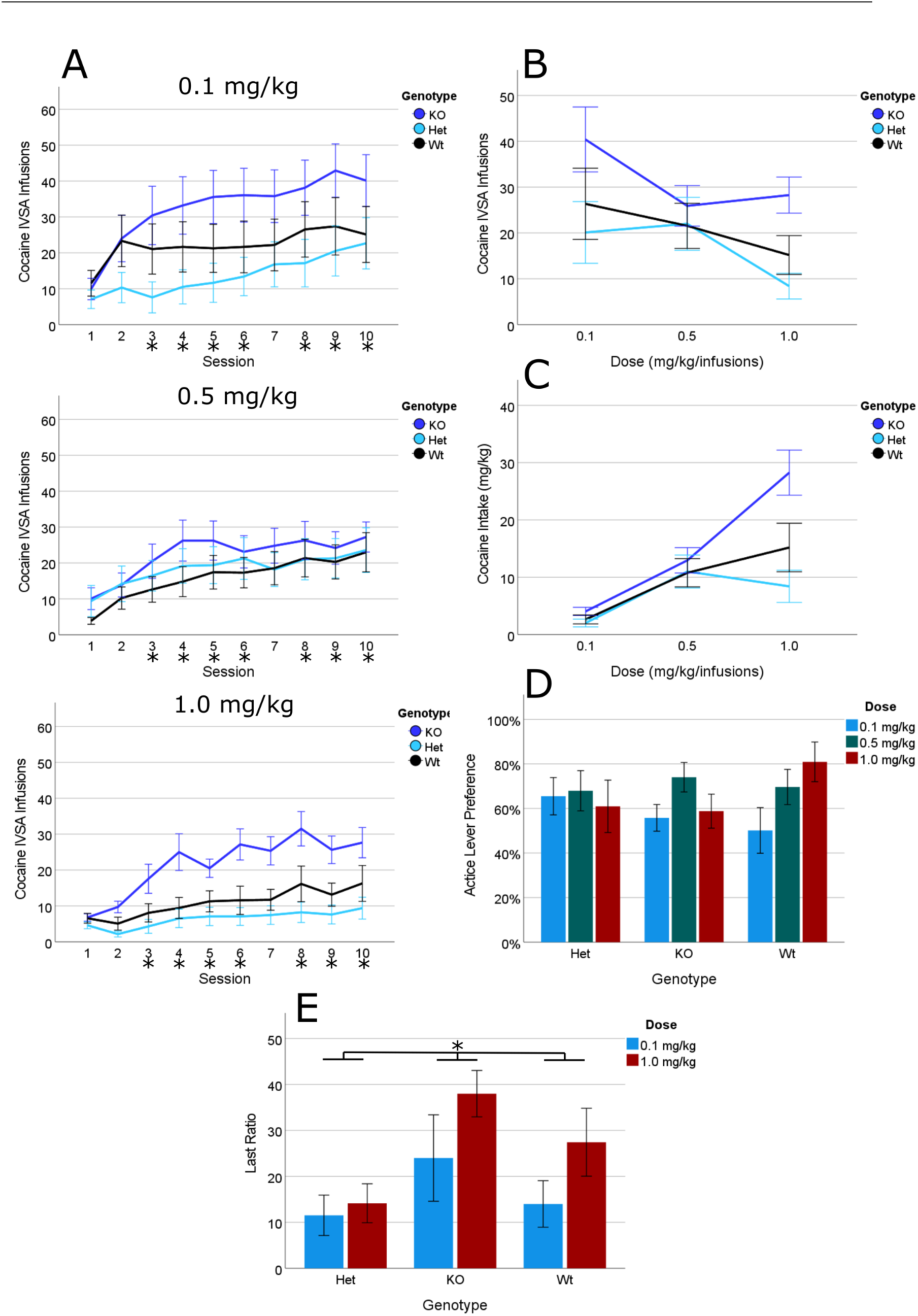
Cocaine IVSA dose responses for wildtype, **HET** and homozygous *Nav1* KO mice. **(A)** 10 sessions of cocaine IVSA for the 3 indicated doses of cocaine using a between-subjects design (n = 13-16 mice per group). *Nav1* KO mice self-administered more cocaine than wildtype and HET mice from session 3 on (* = p<0.05 for post hoc comparisons across dose, no dose by genotype effects were detected). **(B)** A visualization of the dose response curve for the number of cocaine infusions during last 3 sessions, which are averaged for each dose for mice of the indicated genotype. *Nav1* KO mice displayed an upward shift in the dose response curve. **(C)** Visualization of the dose response curve for cocaine intake (mg/kg) during the last 3 sessions for each cocaine dose. Similar to the number of infusions, *Nav1* KO mice displayed an upward shift in the dose response curve for cocaine intake. **(D)** Active lever preference (i.e., the % of correct lever presses) during the last 3 sessions are shown for each of the 3 doses of cocaine. There were no difference in active lever presses among the groups of mice with different genotypes. **(E)** Cocaine IVSA was measured at two different doses (0.1 and 1.0 mg/kg) in Wt, Het and *Nav1* KO mice (n=13-16 mice per group) using a progressive ratio test, where the number of active lever presses required for receiving the next cocaine infusion is progressively doubled from that required for the previous infusion. The number of lever presses required to earn the last cocaine infusion (i.e., the last ratio) provides an assessment of their motivaton for cocaine consumption. *Nav1* KO mice achieved a significantly greater last ratio than wildtype or Het mice at both cocaine doses tested (* p<0.05).

Analysis of cocaine intake also indicated that *Nav1* KO mice self-administered a larger amount of cocaine than HET and wildtype mice at all doses (session-genotype interaction [F(7.7,434.5)=3.0,p=0.003], and there was a main effect of session [F(3.8,434.5)=7.5,p<0.001], and genotype [F(2,113)=9.5,p<0.001]). The pairwise comparisons of the cocaine IVSA results obtained at each session for each genotype indicated that *Nav1* KO mice self-administered more cocaine than HET mice after session 1 and more than wildtype mice after session 2 (p<0.05). Mice of all three genotypes self-administered more cocaine as the dose increased (i.e. there was a main effect of dose [F(2,113)=10.3,p<0.001]) (**Fig. 2C**). Additionally, a genotype-sex interaction [F(2,113)=3.5,p=0.034] suggested that the sex differences were dependent on genotype. Followup of this interaction indicated that there were significant sex differences (males > females) in wildtype (p<0.001) but not in *Nav1* KO (p=0.912) or HET (p=0.218) mice. Similarly, when the preference for the active lever was assessed during the last 3 days of cocaine IVSA, a genotype-sex interaction was detected [F(2,99)=3.3,p=0.041] where wildtype female mice had a lower preference than males (p=0.017) (**Fig. 2D**).

We then assessed their motivation for self-administering cocaine using a progressive ratio schedule of reinforcement (0.1 and 1.0 mg/kg doses), where the number of active lever presses required for receiving the next cocaine infusion was progressively doubled from the response requirement for the previous infusion. The number of lever presses required to earn the last cocaine infusion (i.e., the last ratio) provides an assessment of the motivaton for cocaine consumption. Analysis of the last ratio indicated that there was a genotype-dose-sex [F(2,73)=3.8,p=0.029] and genotype-sex [F(2,73)=8.3,p=0.001] interaction, and a main effect of genotype [F(2,73)=3.9,p=0.025]. Analysis within sex revealed that there was a genotype-dose interaction in males [F(2,38)=4.7,p=0.015] but not in females [F(2,35)=1.3,p=0.292]. Males of the wildtype genotype earned a higher last ratio under the 1 mg/kg dose relative to the 0.1 mg/kg dose (p=0.043). The dose effect in wildtype males appears to be the cause of the genotype-dose-sex interaction. Overall, these data indicate that *Nav1* KO mice maintained their greater level of cocaine intake under a progressive ratio schedule (**Fig. 2E**). In contrast, the acute locomotor responses of *Nav1* KO mice after experimenter administered cocaine was not different from those of C57BL/6 mice (**Fig. S5**).

### The Nav1 KO also impacts food self-administration (FSA)

To determine whether the *Nav1* KO had an impact on another addiction-related phenotype (food reinforcement), mice were tested using a FSA procedure that was similar to that of cocaine IVSA testing except that a chocolate solution was delivered as the reinforcer. Analysis of the number of food reinforcers earned indicated that there was a genotype-session interaction [F(5.1,66.3)=2.5,p=0.036] and a main effect of genotype [F(2,26)=6.5,p=0.005] and session [F(2.6,234)=26.2,p<0.001] (**Fig 3A**). The pairwise comparisons examining the effect of the different genotypes on the FSA results within each session revealed that *Nav1* KO mice earned a greater number of food reinforcers than HET (from session 3 on) or wildtype (from session 4 on except for session 8) (p=0.059) mice (**Fig 3**A). Assessment of active lever preference over the last 3 sessions did not indicate any significant genotypic effect (**Fig. 3B**). *Nav1* KO mice lost more weight than HET or wildtype mice during the period of food restriction, which may be due to their baseline increase in locomotor activity. Therefore, the FSA test was repeated without food restriction. These results confirmed that *Nav1* KO mice earned more food reinforcers: there was a session-genotype interaction [F(5.5,35.6)=3.0, p=0.02], and there was a main effect of genotype [F(2,13)=4.0, p=0.045]) on the FSA results. These results indicate that *Nav1* KO effect on FSA was not due to food-deprivation. After 10 sessions on an fixed ratio1 (FR1) schedule, the mice underwent a progressive ratio test. Analysis of the last ratio achieved also indicated that there was a main effect of genotype [F(2,27)=8.1,p=0.002]. Post hoc, pairwise comparisons indicated that *Nav1* KO mice achieved a greater last ratio than wildtype (p=0.026) and Het (p=0.001) mice (**Fig. 3C**). Taken together, the FSA results indicate that the *Nav1* KO also altered food reinforcement.

**Figure 3.**
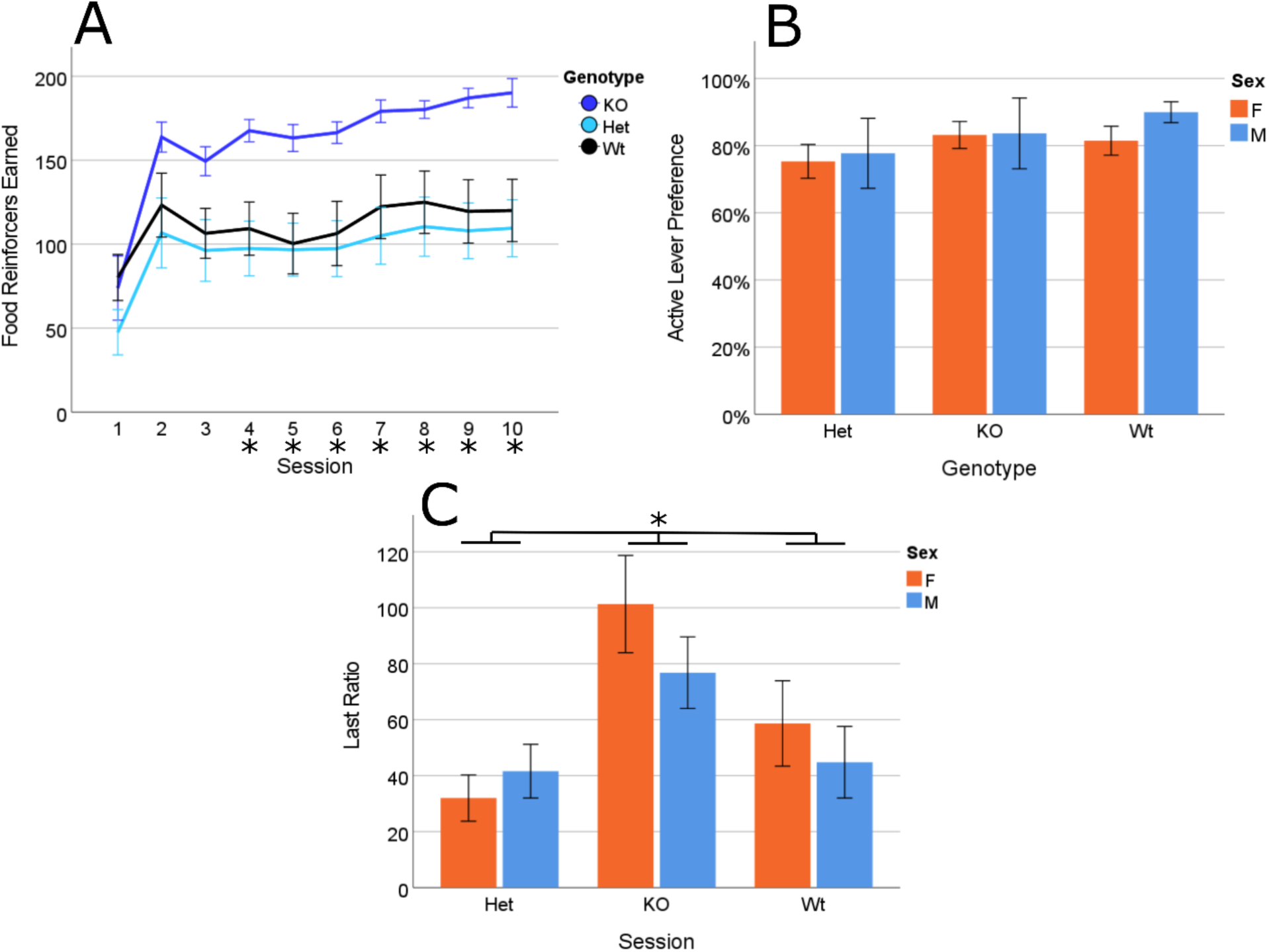
Food self-administration (FSA) by wildtype, HET and *Nav1* KO mice (n=11 mice per group) was measured in 10 daily sessions. **(A)** *Nav1* KO mice had a continuously higher level of FSA than wildtype or HET mice beginning from session 4 (* = p<0.05). **(B)** The percentage of active lever presses (Active Lever Preference) was measured during the last 3 sessions. No differences in active lever preference was detected between mice with the three different genotypes. **(C)** FSA was measured in male and female Wt, Het and *Nav1* KO mice (n=5-6 mice of each sex per group) using a progressive ratio test where the number of active lever presses required for receiving the next food reinforcer is progressively doubled from that required for the previous one. The number of lever presses required to earn the last food reinforcer (i.e., the last ratio) provides an assessment of their motivation for food consumption. Male and female *Nav1* KO mice achieved a significantly greater last ratio than Het or Wt mice (p<0.5).

### Opioid responses were reduced in Nav1 KO mice

Since it was possible that Nav1 could impact responses to other addictive drugs, we examined the *Nav1* KO effect using two murine opioid response assays that model various aspects of the behavior exhibited by human opioid addicts. The morphine conditioned place preference (**mCPP**) test models in mice the drug seeking behavior observed when abstaining humans with an opioid use disorder are confronted with environmental stimuli associated with their drug-taking behavior (*31-34*). In the mCPP assay, after morphine administration is paired with a particular spatial environment, a mouse’s preference for that environment is measured to evaluate the rewarding properties of morphine. During the conditioning phase, C57BL/6J, HET and *Nav1* KO mice exhibited locomotor activation after the morphine injection (p<0.0001), which occurred irrespective of the presence of the *Nav1* KO (p=0.53) (**Fig. 4A**). However, 24 hours after the last conditioning session, C57BL/6J controls (p-value = 0.02) and HET (p-value = 0.033) mice exhibited a preference for the morphine-conditioned side (vs control), while homozygous *Nav1* KO mice did not (p-value > 0.99) (**Fig. 4B**).

**Figure 4.**
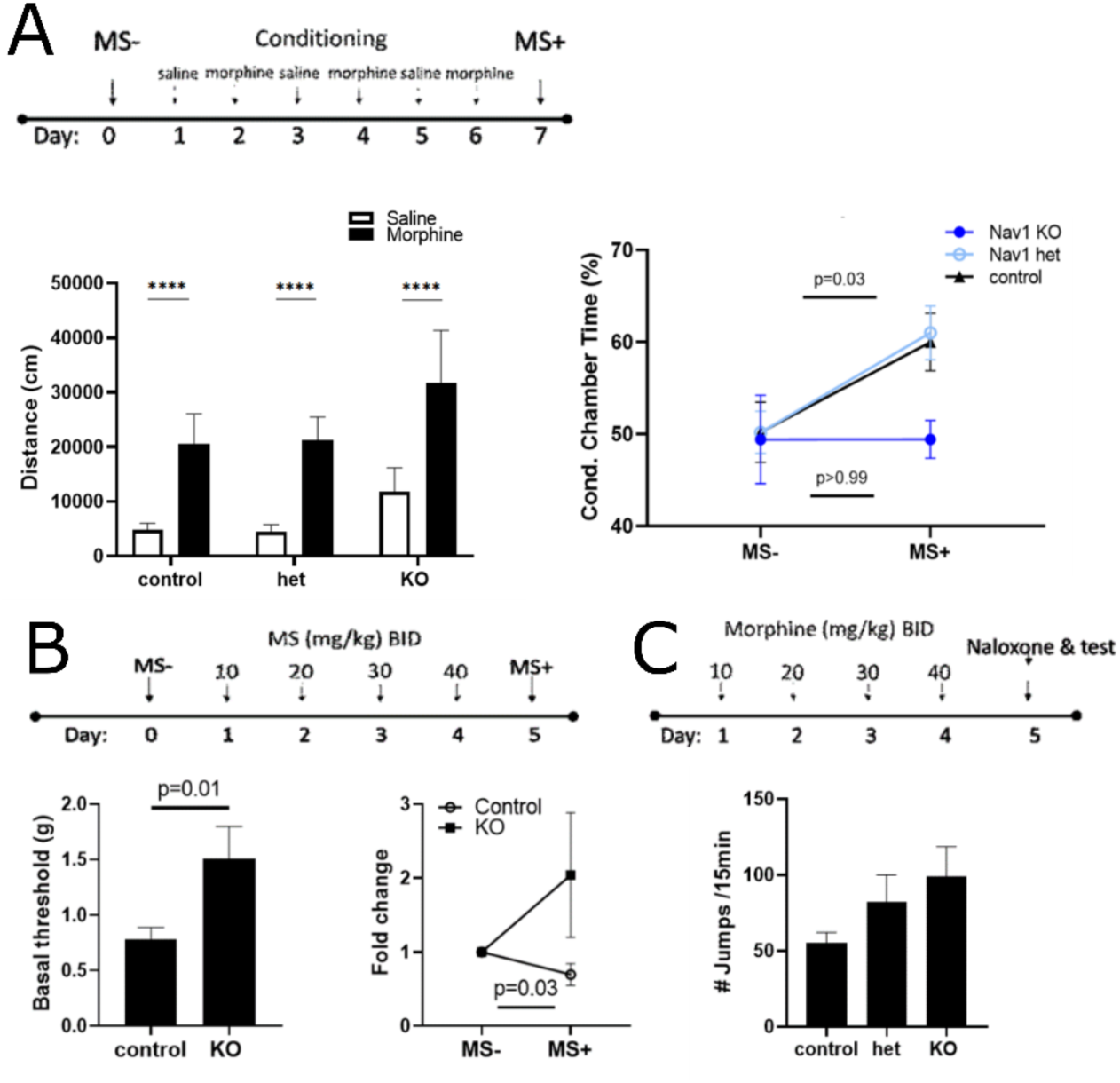
*Nav1* KO mice exhibit altered responses to opiates. **(A)** *Nav1* KO mice maintain their acute locomotor response to morphine, while morphine-induced conditioned place preference (mCPP) is disrupted. The protocol for morphine administration and for mCPP assessment before (M^-^) and after (M^+^) the conditioning sessions where C57BL/6J (control) or *Nav1* KO mice received saline or morphine (5 mg/kg SQ) injections is shown. *Left graph*: The average distance travelled after C57BL/6J, HET or homozygous *Nav1* KO mice received their morphine dose during a one-hour period. ****: p-value<0.0001. *Right:* The time spent in the morphine sulphate administration chamber was measured before (MS-) or after (MS+) undergoing the morphine conditioning regimen (control n=10, Nav1 het KO n=7, Nav1 KO n=7). The p-values assessing the differences between the MS- and MS+ measurements for each genotype are shown. Each bar represents the mean ± SEM of measurements made on 4 – 10 mice per group. **(B)** Tactile sensitivity and opiate-induced hyperalgesia (OIH) are disrupted in *Nav1* KO mice. A timeline for morphine administration and for measuring paw withdrawal is shown. *Left:* At baseline (prior to morphine administration) *Nav1* KO mice have a decreased level of tactile sensitivity (p=0.01) relative to control C57BL/6J mice. Each bar represents the mean ± SEM of measurements made on 12 control and 6 *Nav1* KO mice. *Right:* When tactile sensitivity was measured during the period of opiate withdrawal, C57BL/6J mice have increased tactile sensitivity (i.e., they experience OIH) (P=0.03), while *Nav1* KO mice exhibit a paradoxically reduced level of tactile sensitivity. The fold change is the ratio of the MS+/MS- measurements that were made on 12 C57BL/6J and 6 *Nav1* KO mice. **(C)** *Nav1* KO mice exhibit naloxone-precipitated opiate withdrawal (NPOW) symptoms. A timeline for morphine administration and for the measurement of jumping behavior during NPOW among mice with each of the three genotypes. Each bar represents the mean ± SEM of measurements made on 4 – 10 mice per group.

Besides their analgesic action, opioids also induce a paradoxical hypersensitivity to painful stimuli that occurs during opioid withdrawal in humans (i.e., opiate-induced hyperalgesia, **OIH)**; and inbred strains exhibit large and heritable differences in the extent of OIH developing after chronic morphine exposure (*6, 35*). Therefore, we investigated whether the *Nav1* KO affected the development of OIH. Prior to morphine administration, *Nav1* KO mice exhibited a significantly higher basal nociceptive threshold (i.e., decreased basal tactile sensitivity) relative to C57BL/6J mice (p-value = 0.01) (**Fig. 4C**). After serially increasing doses of morphine (10-40 mg/kg, BID) were administered over a 4-day period, their nociceptive thresholds were measured 18 hrs after the last morphine dose (i.e., during morphine withdrawal to assess the extent of OIH). While C57BL/6J mice exhibited the expected decrease in their nociceptive threshold during morphine withdrawal (p=0.03, paired student t-test), *Nav1* KO mice exhibited a paradoxically reduced level of pain sensitivity (i.e., OIH was absent) (Fig. 4C). In contrast, all three groups of mice exhibited similar levels of naloxone-precipitated opiate withdrawal (NPOW) after morphine treatment, and the *Nav1* KO mice may have exhibited a slightly higher level of NPOW symptoms (**Fig. 4D**) (p=0.08, one-way ANOVA with Tukey’s test). Thus, while *Nav1* KO mice exhibited acute responses to opioids (locomotor activation, NPOW) that were similar to that of control mice, opioid responses that require memory of prior opioid administration (mCPP, OIH) are disrupted in *Nav1* KO mice.

### Nav1 KO mice have alterations in learning, memory, and in other behaviors

Since it was postulated that addictive drugs coopt the reward pathways used for learning and memory (*36*), we investigated whether learning and memory were impacted by the *Nav1* KO. To do this, we first assessed whether the ability to recognize a novel object was altered in *Nav1* KO mice. While C57BL/6J mice spent significantly more time exploring a novel object relative to a familiar one, homozygous and HET *Nav1* KO mice spent a similar amount of time exploring the two objects (**Fig. 5A**) (novelty: p=0.0008; genotype × novelty: p=0.45). There were significant differences in the time that C57BL/6J mice spent with novel vs familiar objects (p=0.045), but this difference was not manifested by HET (p = 1.0) homozygous or *Nav1* KO (p-value = 0.87) mice. When spatial learning and memory capabilities were assessed using the Barnes maze test, *Nav1* KO mice had a significantly reduced ability to correctly recognize the escape hole relative to HET (p<0.01) or C57BL/6J (p<0.0001) mice (**Fig. 5B**). There were no significant differences in the number of primary errors made (p=0.36) or in the total distance travelled (p=0.16) between the three groups of mice. We next investigated whether the *Nav1* KO impacted anxiety or locomotor activity. *Nav1 KO* mice exhibited altered exploratory behavior in the elevated plus maze; they spent more time in the open arms relative to HET (p=0.005) or C57BL/6J mice (p=0.006) (**Fig. S6A**). Open field testing revealed that *Nav1* KO mice had a significant increase in the total distance travelled relative to HET mice (p=0.01); and spent a decreased percentage of time in the open field relative to C57BL/6 mice (p=0.04) (**Fig. S6B**). However, *Nav1* KO mice had normal motor coordination in the rotarod test (**Fig. S6C**). These results indicate that *Nav1* plays an important role in learning and memory capabilities, and that it also effects the level of anxiety and exploratory behavior.

**Figure 5.**
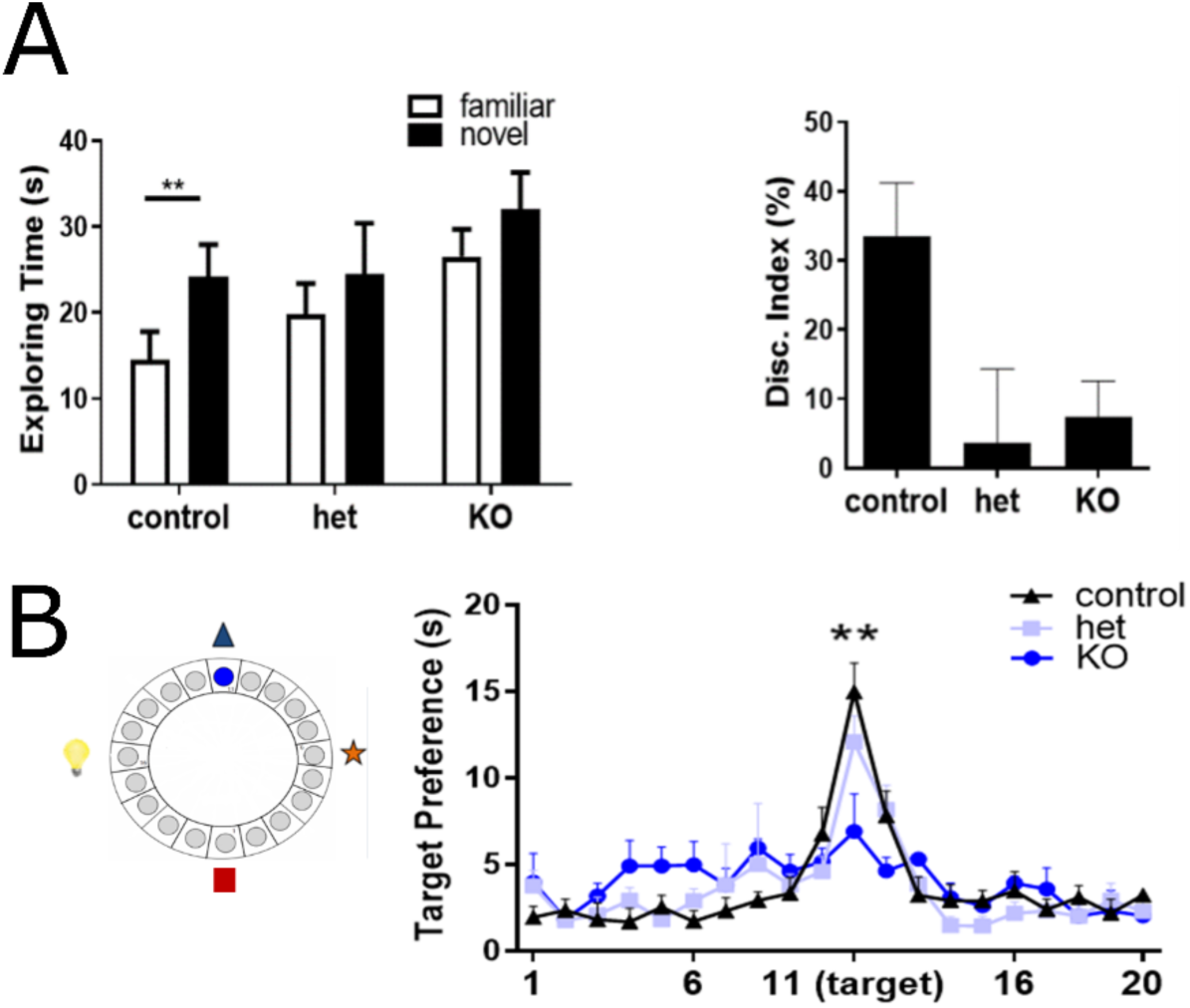
Nav1 KO mice have disrupted novel object recognition and spatial learning abilities. **(A)** The novel object recognition test evaluates their preference for exploring a novel or familiar object. The left graph compares the time spent exploring a novel or familiar object (**, p<0.01). In the right graph, the discrimination index is calculated as the percentage of time spent with (novel – familiar object)/ total exploration time. Each bar represents the mean ± SEM for measurements made on 9 C57BL/6J (control), 7 HET, and 10 *Nav1* KO mice. Control mice spent more time with the novel object (**, p<0.01), while HET and homozygous *Nav1* KO mice did not. (**B**) The Barnes Maze test evaluates spatial learning and memory abilities. A schematic diagram indicates how the ability of a mouse to correctly identify a target escape hole (shown in blue) with the aid of visual cues that are shown outside the circle. The graph shows the time spent at the target hole ± SEM during a 90 second test session, which was measured after 12 training sessions. The *Nav1* KO mice had a significantly reduced ability (** p<0.01) to correctly identify the target hole. Each data point is the average of measurements made on 6 control, 7 HET *Nav1*, and 4 homozygous *Nav1* KO mice.

### Increased excitatory hippocampal synaptic transmission in Nav1 KO mice

Since Nav1 is involved in neuronal development and migration (*22, 23*), we quantitatively examined the impact of the *Nav1* KO on synapse formation. *Nav1* KO mice had a significant increase in the number of excitatory synapses in the dentate gyrus of the hippocampus (vs C57BL/6J mice, p=0.03), but inhibitory synapse density was not changed (p>0.99) (**Fig. 6A**). *In vitro* electrophysiological recordings from granule cells in the dentate gyrus of the hippocampus showed a significant increase in the frequency (but not in the amplitude) of miniature excitatory postsynaptic currents (mEPSCs) in *Nav1* KO vs C57BL/6J mice (**Fig. 6B**), which indicates that there is increased excitatory synaptic transmission in the dentate gyrus of *Nav1* KO mice. In contrast, there was no significant difference in the frequency or amplitude of the miniature inhibitory postsynaptic currents (mIPSCs) measured in *Nav1* KO vs C57BL/6J mice. There was also a significant decrease in the number of inhibitory synapses formed in the prefrontal cortex (**PFC**) of *Nav1* KO mice (vs. C57BL/6J, p=0.008), while excitatory synapse density was not altered (Fig. 6A). Since the nucleus accumbens (**NAc**) is a critical center that mediates responses to addictive drugs (*37*), we compared the cells in the NAc of *Nav1* KO and C57BL/6J mice by immunohistochemical staining. There was a similar density of neuronal cells, as shown by the equal number of neuronal nuclei marker^+^ and neurofilament H^+^ cells. However, glial fibrillary acidic protein (GFAP) staining indicated that there was a slight trend toward an increase in the number of glial cells in the NAc of the *Nav1* KO (**Fig. S7**). Overall, the cellular composition of the NAc appeared to be less affected by the *Nav1* KO than that of the hippocampus and PFC.

**Figure 6.**
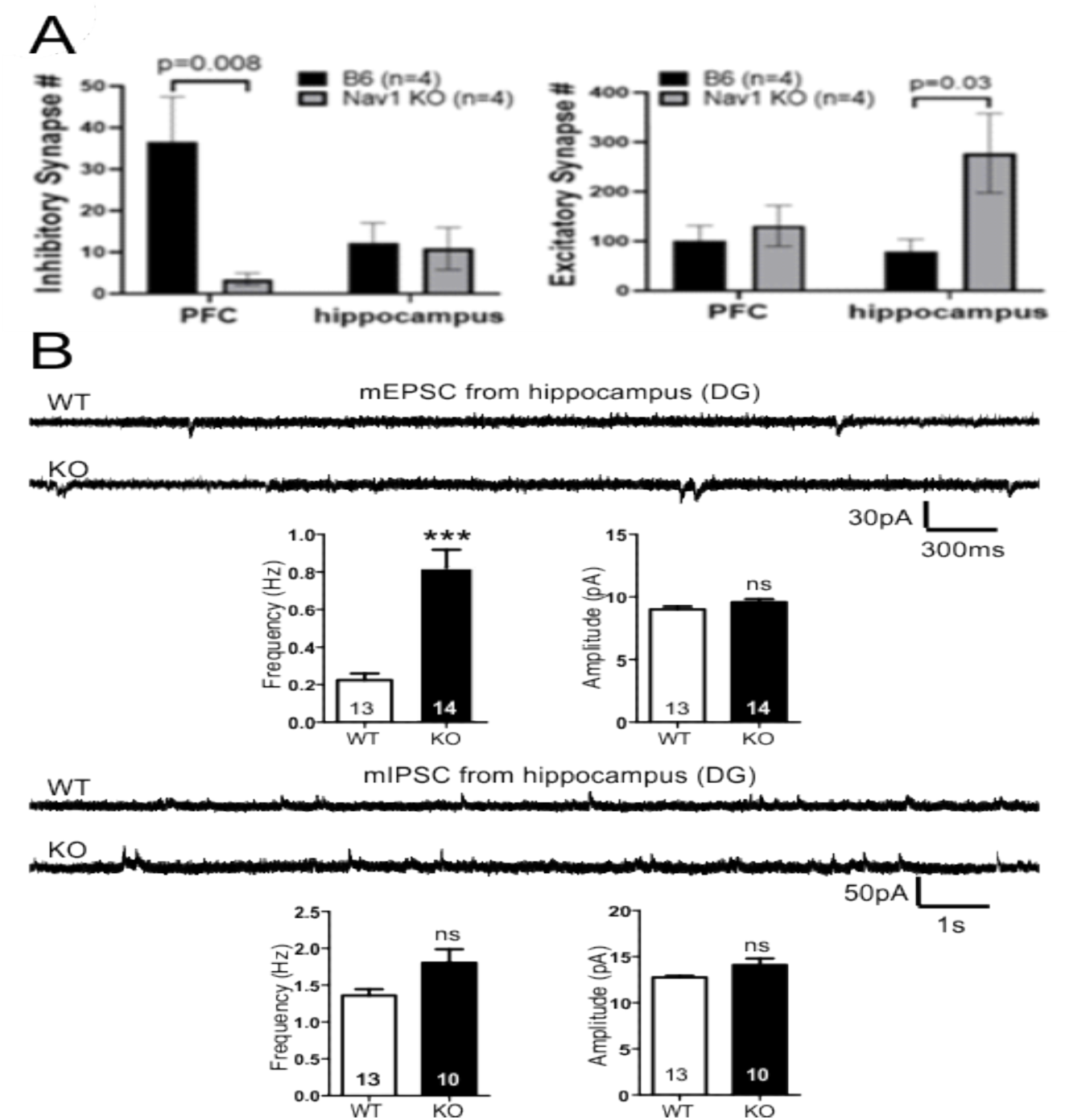
**(A)** Excitatory and inhibitory synapse formation in *Nav1* KO mice is disrupted. Brain slices prepared from hippocampal and PFC of *Nav1* KO mice were immunostained with antibodies to inhibitory presynaptic (vGat) and post-synaptic (gephyrin) marker proteins; or with antibodies to excitatory presynaptic (vGlut1) and post-synaptic (PSD95) marker proteins. Each bar is the average ± SEM of the number of inhibitory or excitatory synapses measured in PFC or hippocampal slices generated from four isogenic C57BL/6J or *Nav1* KO mice. Inhibitory synapse density in the PFC was significantly decreased (p=0.008) while the excitatory synapse density in the hippocampus was increased (p=0.03) in *Nav1* KO mice. (**B**) Excitatory synaptic transmission in the dentate gyrus of the hippocampus of *Nav1* KO mice is increased. Representative traces of mEPSCs (Top) or mIPSCs (Bottom) recorded from granule cells in the dentate gyrus of the hippocampus of C57BL/6J (WT) and *Nav1* KO mice. The graphs show a significant increase in mEPSC frequency in granule cells of *Nav1* KO (vs. wildtype) mice, but not in their amplitude. In contrast, there was no significant difference in mIPSC frequency or amplitude between *Nav1* KO and WT mice. The numbers within the bars indicate the number of cells evaluated. ***P < 0.001 by unpaired t-test. ns: not significant.

These results indicate that *Nav1* KO mice have an increase in hippocampal excitatory synapses and synaptic transmission, and that there is a decrease in the number of inhibitory synapses formed in the PFC.

### Nav1 KO-induced transcriptomic changes

Since the PFC regulates higher cognitive functions (i.e., learning, memory, and decision making) and forms connections with multiple other brain regions (*38, 39*), we examined *Nav1* KO-induced transcriptomic changes in the PFC. snRNA-Seq was performed on 28,686 high quality PFC cells obtained from *Nav1* KO mice (14056 cells; 68637 reads per cell) and from age-matched, isogenic C57BL/6J mice (14630 cells, 56980 reads per cell with a mean of 1438 genes per cell). The PFC cells were separated into 14 clusters, which based upon their pattern of marker mRNA expression (*40*), were derived from 4 lineages: neurons, astrocytes, oligodendrocytes, and microglia (**Figs. 6-7, S8-S9).** The eight neuronal clusters were separated into six excitatory (0-2, 4, 5, and 10) and two (3, 8) inhibitory neuronal clusters. Consistent with prior findings (*40*), excitatory neurons were 5 to 7-fold more abundant than inhibitory neurons (52-75% vs 7-14% of the total). The cell clusters and their distribution in our C57BL/6J PFC data was very similar to that of a prior analysis (*40*) (**Fig. S10**). Clusters 10 and 5 had the highest levels of *Nav1* mRNA (Fig. 8). Cluster 10 was of particular interest since its abundance was most increased (14-fold) in *Nav1* KO (vs C57BL/6J) mice. Cluster 10 uniquely expressed an excitatory neuronal marker (*Tshz2*) that is characteristic of cortical layer 5 (**L5**) cells (Fig. 8), which form projections to extra-cortical brain regions and are in the cortical layer whose transcriptome was most altered during cocaine withdrawal (*40*). Pathway enrichment analysis of the differentially expressed genes (**DEGs**) in cluster 10, revealed that many of the enriched pathways were associated with neuronal guidance, synapse or signaling functions; and 7 DEGs (*Gria3, Grin2b, Grin2a, Camk2a, Ppp3ca, Gria4, Prkcb*) were associated with an amphetamine addiction pathway (p=0.000025) (**Fig. S11)**. Clusters 5 and 10 also had the highest levels of expression of two mRNAs (*Rab3a* (*41*), *Cck* (*42*)), whose expression levels were altered during cocaine withdrawal (*40*) (**Fig. S12**). Cluster 5 uniquely expressed a transcription factor (*Etv1,* aka *Er8*) that is required for generating habitual behaviors (*43*). The inhibitory neurons in cluster 8 were of interest because their abundance was 3-fold decreased in the PFC of *Nav1* KO mice, and they uniquely expressed *Drd1* mRNA (Fig. S11). In contrast, *Drd2* mRNA was not expressed at detectable levels in any cluster. In summary, the transcriptomic analysis indicates that the *Nav1* KO affected neurons located in a deep cortical layer, and some of these cells had transcriptomic changes that were similar to those associated with cocaine withdrawal.

**Figure 7.**
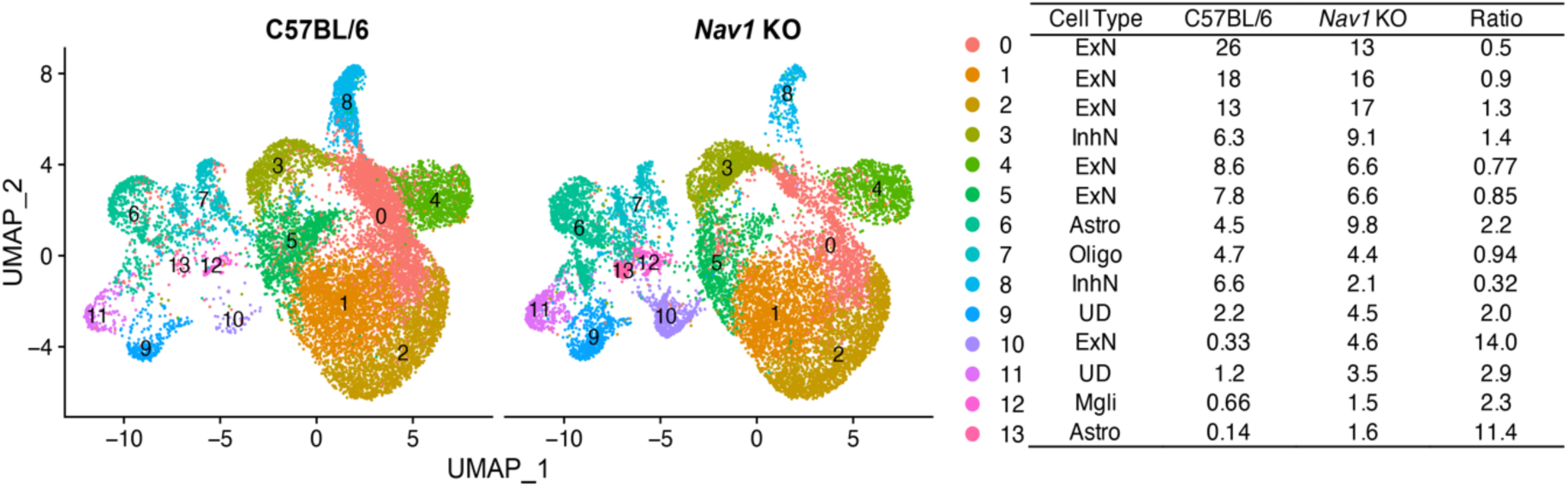
snRNA-Seq analysis of *Nav1* KO and isogenic C57BL/6J PFC tissue. UMAP plots show the data for the C57BL/6J and *Nav1* KO cells. Each dot represents an individual cell. The table indicates the dot color for each cluster, the percentage of cells in each cluster, and the ratio of the cell percentages (*Nav1 KO*/C57BL/6J) in C57BL/6J and *Nav1* KO PFC. This dataset analyzed the expression of 21,892 genes in 28,686 nuclei. Based upon total transcriptomic differences, the PFC cells were separated into 14 different clusters. Based upon canonical marker expression, the clusters are derived from the five lineages: astrocytes (6, 11), microglia (12), oligodendrocytes (7); and inhibitory (3, 8) and excitatory (0, 1, 2, 4, 5, 10) neurons. The lineages for two clusters (9, 11) were undefined (UD) because they expressed markers from different cell lineages. Of particular interest, the number of cells in an inhibitory neuron cluster (cluster 8) is three-fold decreased in *Nav1* KO mice, and the number of cells in an excitatory neuron cluster (cluster 10) is 14-fold increased in *Nav1* KO mice.

**Figure 8.**
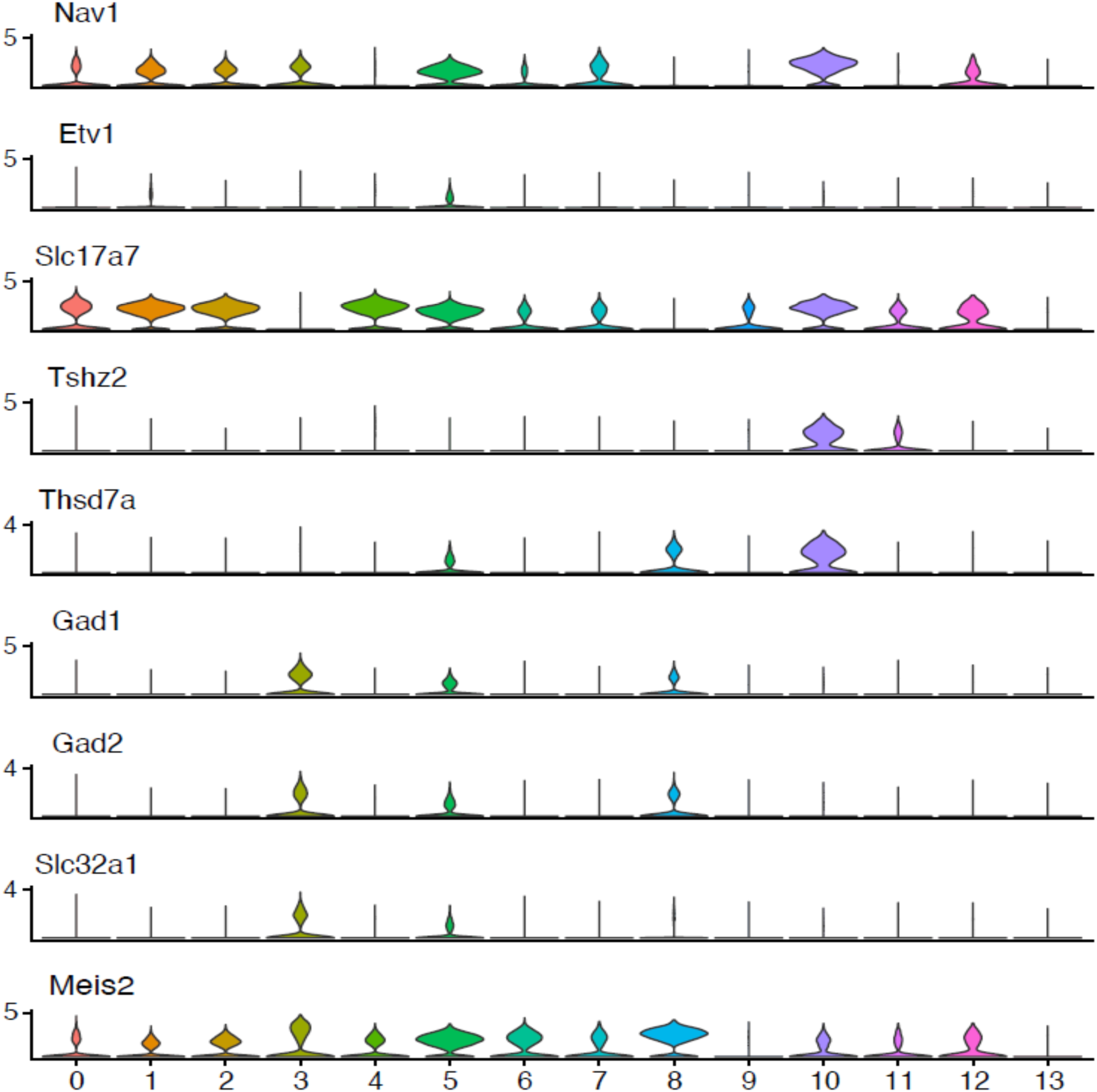
Violin plots showing the level of expression of *Nav1*, *Etv1* and 7 neuronal marker mRNAs within 14 cell clusters identified in the PFC. The cluster number is indicated on the x-axis and the y-axis shows the natural log transformed and normalized level of expression of each mRNA. Neuronal cluster identity was determined by expression of canonical mRNAs (*40*) for excitatory (*Slc17a7, Tshz2, Thsd7a*) and inhibitory (*Gad1, Gad2, Slc32a1, Meis2*) neurons. Cluster 10 was the only cluster with a defined lineage that expressed a marker (*Tshz2)* for layer 5 cells, which are the corctical neuron whose transcriptome was altered during cocaine withdrawal (*40*). Cluster 11 identity could not be determined due to expression of mRNAs for different lineages. Cluster 5 cells uniquely expressed mRNA for a transcription factor (*Etv1*) required for habitual behavior (*43*).

## Discussion

Our analysis of a murine genetic model identified *Nav1* as a candidate gene that influences voluntary cocaine consumption, and this genetic finding was confirmed by the increased level of cocaine IVSA exhibited by *Nav1* KO mice across a broad range of cocaine doses. The effect of the *Nav1* KO on voluntary cocaine consumption occurred early; and it was maintained throughout a 10-day period of testing and during subsequent progressive ratio testing, which tests their motivation for consuming cocaine by substantially increasing the effort required to receive a dose of cocaine (*44*). Since *Nav1* KO and wildtype mice did not exhibit differences in lever preference or in cocaine-induced acute locomotor responses; neither indiscriminate lever pressing nor differential locomotor effects could be responsible for their increased cocaine-intake. Overall, these results indicate that the reinforcer efficacy of cocaine is increased in *Nav1* KO mice, which produces an increased motivation to consume cocaine. The *Nav1* KO had a similar effect on food reinforcement, which suggests that the *Nav1* KO effect could be generalized across drug and non-drug reinforcers. It has been postulated that a common neural circuitry may underlie food and drug addictions (*45*), and genetic variation in *Cyfip1* had effects on both cocaine and food reward (*5, 46*). *Nav1* KO mice also exhibited reduced anxiety-like behavior. An inverse relationship between anxiety-like behavior and cocaine IVSA (*47*) and ethanol drinking (*48*) was previously observed in rats. While the Nav1 results also indicate that the neural circuits mediating drug reward, food reward and anxiety may have some overlap, additional research is necessary to explore this possibility further.

The Nav1 effect extended beyond cocaine and food reinforcement to opioid responses. While the *Nav1* KO increased cocaine reinforcer efficacy in the IVSA procedure, it blunted the opioid response in two assays (CPP and OIH). The CPP and IVSA results obtained for different drug types are not always the same (*49, 50*) nor are the results always consistent (*34*). The previously observed inverse effect of a genetic factor on cocaine CPP and IVSA results suggests that a common genetic factor may exert opposite directional effects on these two measures (*49*). While spatial learning may be required for CPP (*51*) to be manifest after chronic opioid exposure, *Nav1* KO mice did not develop OIH after chronic morphine administration. Since learning is not required for OIH to develop, this confirms that Nav1’s blunting of opioid responses was not due to a learning defect. Our own analyses of murine genetic models indicate that the appearance of OIH after chronic opioid exposure is under genetic control (*6, 7, 35, 52*). The mechanism by which *Nav1* affects OIH is not known, but the *Nav1* KO-induced change in the basal nociceptive threshold may provide a clue about its effector mechanism. Collectively, our results indicate that *Nav1* alters the reward response to both opioids and cocaine.

*Nav1* KO mice also exhibited reduced spatial learning and memory. The *Nav1* KO induced alterations in the balance between excitatory and inhibitory synapses in the cortex and hippocampus provides a potential mechanism by which *Nav1* could impact learning, memory, and the responses to addictive drugs. This mechanism is consistent with the known role of *Nav1* in regulating neuronal development, directional migration (*22*), and neurite outgrowth (*23*). While various cortical and subcortical brain regions affect the response to addictive drugs, the PFC integrates reward-seeking and decision-making, through the inhibitory control it exerts (*14, 53*) through the dense reciprocal connections it forms with virtually all neuromodulatory centers within cortical and subcortical regions (*54*). For example, PFC glutamatergic neurons connect to the NAc core, which can provide a top-down control mechanism for preventing food addiction behaviors (*55*). Disruptions to hippocampal synaptic balance could be a cause of the learning/memory deficits observed in *Nav1* KO mice, since these tests (particularly the Barnes maze) are hippocampal-dependent (*56, 57*). Since the hippocampus also has a role in cocaine responses (*58*), *Nav1-*induced disruptions of hippocampal function could also affect cocaine IVSA.

In summary, we demonstrated that *Nav1* impacts voluntary cocaine consumption, food reinforcement, opioid reward and nociceptive responses, anxiety-like behavior, learning/memory, and the excitatory/inhibitory synaptic balance in hippocampal and PFC brain regions. Collectively, these results highlight the important role that Nav1 plays in multiple addiction-related traits. Further characterization of neural adaptations occurring in *Nav1* KO mice could enable the pathways mediating addiction-associated behaviors to be more fully characterized.

## Abbreviations

CPP: conditioned place preference
DEGs: differentially expressed genes
cSNPs: coding single nucleotide polymorphism
HBCGM: aplotype-based computational genetic mapping
HMDP: hybrid mouse diversity panel
IVSA: intravenous self-administration
KO: knockout
mEPSCs: miniature excitatory postsynaptic currents
mIPSCs: miniature inhibitory postsynaptic currents
MRI: magnetic resonance imaging
NPOW: naloxone-precipitated opiate withdrawal
OIH: opiate induced hyperalgesia
PFC: prefrontal cortex
SUD: substance use disorder

## Funding

This work was supported by a NIH/NIDA award (5U01DA04439902) to GP and T32AA0256606 (JDJ, JRB).

## Data availability

All raw single cell RNA-seq data and the processed data were deposited in the Gene Expression Omnibus (GEO): GSE216957. SNP allele data is also available at the Mouse Phenome Database (GenomeMUSter https://mpd.jax.org/genotypes).

## Author contributions

GP and JDJ formulated the project. YT, JRB, JDJ and GP wrote the paper with input from all authors. JRB, YT, WZ, ZC, ST, YG, LJ, RJ, DP, FG, and IP generated experimental data. JRB, ZF, AA, and MW analyzed the data. All authors have read and approved of the manuscript.

## Supplemental Information

### Materials and Methods

#### Mouse Strains

Twenty-one inbred strains (see Fig 1A for strains) and *Nav1* knock out (KO) mice (on C57BL/6J background) were maintained at Binghamton University and at the Stanford University School of Medicine. All mice were housed in their home cage and maintained on *ad libitum* mouse chow (5L0D, Purina Lab Diet) and water. Mice were individually housed in polycarbonate cages (30 × 8 cm) with wood-chip bedding (SANI-CHIPS), a paper nestlet and a red polycarbonate hut. All procedures were approved by the Binghamton University Institutional Animal Care and Use Committee or by the Stanford Institutional Animal Care and Use Committee; and were conducted in accordance with the National Institute of Health “Guide for Care and Use of Laboratory Animals, Eighth Edition” (National Research Council (US) Committee for the Update of the Guide for the Care and Use of Laboratory Animals, 2011). All mice were originally obtained from Jackson Laboratories, and the results are reported according to the ARRIVE guidelines (*1*).

#### Cocaine intravenous self-administration (IVSA)

Mice were anesthetized with isoflurane (3% at induction, 1.5-2% for maintenance) and implanted with a chronic indwelling jugular catheter (catalog # CNC-2/3S-082109E/12, Access Technology, IL USA) and access button port (1-VAB62SMBS/25, Instech, PA USA). Carprofen (5 mg/kg) was administered by subcutaneous injection pre- and post-surgically (10-14 hrs from the previous dose), for a total of 5 doses. 1 mL warm saline was administered by subcutaneous injection upon completing the surgery and again 24 hours later. Catheters were maintained with a flush of ∼0.05 mL of sterile saline and then filling the catheter with heparin lock solution (∼0.01 mL volume, 500 units/mL concentration, SAI Infusion Technologies, IL) at least once every 3 days during recovery periods and daily during IVSA testing. Catheter patency was confirmed by infusions of propofol (∼0.02 mL volume, 10 mg/mL concentration, Zoetis, NJ). Immediate but rapidly reversed loss of muscle tone indicated that the catheter was patent. This testing occurred once before the start of IVSA testing (3-4 days prior to testing) and again after the mouse completed the final testing session. Any mouse that failed the test was excluded from the experiment.

Twenty-one strains (n = 1-6 mice per strain, all male) were tested for cocaine IVSA acquisition in 10 consecutive daily sessions (Fixed-ratio-1 schedule of reinforcement, 0 or 0.5 mg/kg body weight of cocaine per infusion) that ran until 65 infusions were earned or 2 h passed, whichever came first. Cocaine hydrochloride (Sigma Aldrich; St Louis MO) was dissolved in sterile saline at a concentration of 0.17, 0.84, or 1.68 mg/mL to produce a freebase dose of 0.1, 0.5, or 1.0 mg/kg/infusion (infusion volume was 0.67 mL/kg/infusion). Testing occurred at the same time each day, during the light phase of a 12/12 h cycle. The animals were tested in Med Associates mouse self-administration chambers (55.69 x 38.1 x 35.56 cm, MED-307W-CT-D1, Med Associates, VT) that were fitted with 2 retractable ultrasensitive levers and that were housed within sound-attenuating cubicles. Assignment of the active infusion lever (right or left side of the box) was counterbalanced across strains/sex. Assignment of testing chamber minimized testing multiple mice from a given strain in the same chamber. Test sessions began with the activation of the white noise and the illumination of 5 stimulus lights on the back wall of the chamber. No priming infusion(s) were delivered. When a subject actuated the active lever, an infusion was delivered, the house light flashed, and the aperture lights turned off for 20 s. During this time-out period, contacts on the active lever were recorded but had no programmed consequence. Actuation of the inactive lever had no programmed consequence.

The number of infusions earned, cocaine intake (mg/kg) (number of infusions X dose) and active lever preference (active lever presses/total lever presses) were key variables of interest. Active lever preference is incalculable if the mouse fails to press either lever, and consequently analysis of this variable by session leads to excessive missing data points. Therefore, the last 3 IVSA sessions were collapsed by calculating preference across these days, within individual mice.

#### Haplotype based computational genetic mapping (HBCGM)

The SNP database was generated by analysis of the genomic sequences of 47 classical inbred strains (C57BL/6J, 129P2, 129S1, 129S5, AKR, A_J, B10, BPL, BPN, BTBR, BUB, BALB, C3H, C57BL10J, C57BL6NJ, C57BRcd, C57LJ, C58, CBA, CEJ, DBA, DBA1J, FVB, ILNJ, KK, LGJ, LPJ, MAMy, NOD, NON, NOR, NUJ, NZB, NZO, NZW, PJ, PLJ, RFJ, RHJ, RIIIS, SEA, SJL, SMJ, ST, SWR, TALLYHO, RBF, MRL) and of 6 wild-derived inbred strains (CAST, MOLF, PWD, PWK, SPRET, WSB) as described in (*2-4*). HBCGM was performed as originally described (*5*) using modifications described in (*6*). In brief, HBCGM utilizes bi-allelic SNPs that are polymorphic among the strains. Only bi-allelic SNPs with at least one definitive homozygous alternative call among the strains were included, while all other variants (including INDELs) were marked and excluded from HT-HBCGM analysis. The chromosomal location and potential codon-change caused by a SNP are then annotated relative to the encoded gene using predictive gene models from Ensembl version 65. Only SNPs meeting the following criteria were used for haplotype block construction: (i) polymorphic among the strains with input trait data; and (ii) there were at least 8 strains with unambiguous allele calls, which is an important criterion because it ensures that there is sufficient genetic diversity in the analyzed cohort for analysis by HBCGM. Haplotype blocks with 2, 3, 4 or 5 haplotypes were then dynamically produced and the correlation between the input phenotypic data and the haplotype pattern within each identified block was evaluated as described (*6*). Next, the genes are then sorted based upon the ANOVA p value (in increasing order) for numeric data or by the F statistic (in decreasing order) for categorical data. For a gene that is covered by multiple blocks, the smallest p value (or largest F statistic) obtained for all blocks within that gene was used. The genetic effect size (η^2^) is calculated as previously described (*7, 8*):

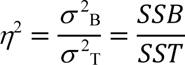

where SSB is the between-group sum-of-squares of the ANOVA model and SST is the total sum-of-squares. η^2^ is the genetic effect of the groups defined by haplotypes on the trait value and the total variance (σ^2^T) consists of within-group variance and between-group variance as:

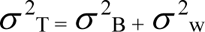

for a sample size of n with *k* groups, with equal group sizes the *F* statistics of samples with effect size *η*^2^ follows a noncentral *F* distribution as *F*(k – 1, n – k, λ) with the non-centrality parameter:

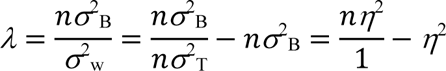

Therefore, the significance level α for power of one-way ANOVA test is given as:

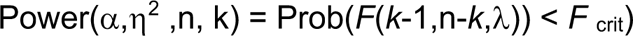

where *F*_crit_ = *F*_(1−α, k–1, n–*k*)_ is the (1−α) quantile of the *F* distribution with *k* – 1 and n – *k* degrees of freedom. Additional details about the HBCGM method are described elsewhere (*8*).

### Generation of Nav1 KO mice

C57BL/6 female mice were super-ovulated by intraperitoneal injection of pregnant mare’s serum gonadotropin and human chorionic gonadotropin. These mice were then paired with C57BL/6 males to generate fertilized embryos, and pronucleus (PN) stage embryos were collected. Cas9 and 2 sgRNAs (CAAACCTAGCCGGATTCCTC, GCACGGTAACCACAAGCTCG) were then electroporated into PN embryos using a NEPA21 electroporation system (Fig. S2). The sgRNAs were designed to delete a 178 bp region at the end of exon 1 and to also introduce an early stop codon in exon 1. This region was deleted because exon 1 is expressed in all 7 known isoforms of *Nav1* mRNAs. Healthy embryos were transferred into the oviducts of pseudo-pregnant recipient females. Genomic DNA from the pups were screened by PCR amplification using the strategy shown in Fig. S2. Mice with genomic DNA that generated diagnostic amplicons were subsequently sequenced to characterize the deleted region.

### smFISH analysis

Single molecule fluorescence in situ hybridization (smFISH) was performed according to (*9*). In brief, frozen brain tissue sections (20 µm) were pre-treated with 0.01% pepsin in 0.1 M HCl for 2 min at room temperature followed by washing in 0.05% Tween 20 in 1 x diethyl pyrocarbonate-treated (DEPC)-PBS. mRNA was reverse transcribed to cDNA in a room temperature buffer containing 0.5 mM dNTP, 0.2 μg/μl BSA, 1 μM cDNA primer, 1 U/μl RNaseIn (Clonetech, 2313B) and 20 U/μl RT (Maxima, Thermo Scientific™, EP0752) for 3 hr at 50 °C in a securely sealed chamber. After three brief washes in 0.05% Tween 20 in 1 x DEPC-PBS, the sections were post-fixed in 4% paraformaldehyde for 30 min at room temperature and washed three times in phosphate-buffered saline with Tween 20 (PBST). Hybridization and ligation with T4 DNA Ligase were performed in T4 ligase buffer with 0.2 μg/μl BSA, 100nM padlock probe and 0.1 U/μl T4 DNA Ligase (New England Biolabs) for 30 - 45 min in 37 °C. This was followed by washing with 2× SSC with 0.05% Tween-20 at 37 °C for 5 min, and then rinsing in PBST. Rolling circle amplification (RCA) was performed with 1 U/μl Φ29 DNA polymerase (New England Biolabs) using the reaction buffer supplied by the manufacturer with 250 μM dNTPs, 0.2 μg/μl BSA and 10% glycerol. The incubation was carried out for 60 - 150 min at 30°C, which was followed by a washing in TBST. Stranded RCA products (RCP) were hybridized in 250 nM of Cy5 and FAM fluorescence-labeled oligonucleotide probes in a solution of 2X SSC, 20% formamide for 30 min at 37 °C. Slides were then washed in TBST. Dried slides were mounted with VECTASHIELD® PLUS Antifade Mounting Medium (vectorlabs, H-1900-10). Images were acquired using an SP8 Confocal microscope (Leica). For quantification, the number of RCPs and cell nuclei in the images were counted digitally using Fiji software (version 1.53C) (*10*). All oligos listed in Table S4 were synthesized by Integrated DNA Technologies, Inc (Coralville, Iowa).

### Proteomic mapping

Proteins in brain tissue obtained from C57BL/6 and Nav1 KO (N93) mice were extracted and separated by SDS-PAGE. Protein bands corresponding with the molecular weight of Nav1 were excised, and trypsin digested. The digested peptide mixtures were z-tip purified and run on an Orbitrap Fusion™ Lumos™ Tribrid™ Mass Spectrometer (ThermoFisher, San Jose), which was equipped with an Acquity UPLC M-class system (Waters, MA). The peptide data were searched against the mouse proteome database. Two different Nav1 peptides were found in the C57BL/6 brain sample (MW 202.423; calculated pI 8.06, Score Sequest HT 2.00323, n=2 peptides; and MW 252.876, calculated pI 8.76, Score Sequest 2.131383, n=1 peptide). In contrast Nav1 peptides were completely absent in brain tissue obtained from the *Nav1* KO. Thus, proteomic mapping confirmed that Nav1 protein was absent from brain tissue obtained from homozygous *Nav1* KO mice, while Nav1 peptides were detected in C57BL/6 brain tissue.

### Cocaine IVSA phenotyping of Nav1 KO mice

C57BL/6J (Wt), heterozygous *Nav1* KO (Het) and homozygous *Nav1* KO mice (KO), which were all naïve to any prior experimentation, were tested in a between-subjects dose-response test using the IVSA procedures described above with cocaine doses of 0.1, 0.5, and 1.0 mg/kg. The following numbers of mice of the indicated sex were tested at each of the indicated doses: **KO** – 0.1 Dose: 7F, 7M; 0.5 Dose: 8F, 8M; 1.0 Dose: 6F, 8M; **Het** – 0.1 Dose: 7F, 8M; 0.5 Dose: 7F, 8M; 1.0 Dose: 6F, 7M; and **Wt** – 0.1 Dose: 7F, 8M; 0.5 Dose: 7F, 9M; 1.0 Dose: 8F, 6M.

#### Food Self-Administration (FSA) testing

Mice were tested in a FSA procedure, which utilized the same testing conditions as cocaine IVSA, except that 20 µl of Chocolate Boost (Nestle) was delivered as the reinforcer. Additionally, the number of reinforcers per session was not limited to avoid any ceiling effects; the mice had no prior surgical procedures and were not tethered to an infusion line. All sessions were terminated after 2 hours of testing. To facilitate acquisition of FSA, home-cage food was removed ∼16 hours before the 1^st^ FSA session. Following the 1^st^ session, the mice were fed 2.5 grams of food in the home-cage, in order to maintain food restriction through the 2^nd^ session. Following the 2^nd^ session, mice were returned to *ad lib* feeding for the remaining of the testing period. The following numbers of mice of the indicated sex were tested in the FSA assay: **KO** – 6F, 5M; **Het** – 6F, 5M; and **Wt** – 6F, 5M.

Following the 1^st^ FSA experiment, we noted that homozygous *Nav1* KO mice lost more body weight, relative to HET and wt mice, in response to food deprivation. Therefore, we tested a second cohort of mice under no food restriction. This cohort was instead subjected to magazine training prior to FSA sessions. This training involved placing mice in the operant chambers and activating a program that turned on the white noise and illuminated the nose-poke apertures and house light. After 10 seconds, 20 µl of Boost was delivered to the magazine, the aperture lights turned off and the house light flashed (1 sec on, 1 sec off). These conditions continued until the mouse entered the magazine, as detected by infrared beam break. Entry to the magazine stopped the flashing house light and illuminated the apertures. 30 seconds following the magazine entry, 20 µl of Boost was again delivered to the magazine, the aperture lights turned off and the house light flashed until another magazine entry. The sessions were limited to 50 Boost rewards or 2 hours. Mice were required to earn at least 30 rewards before moving on to FSA testing.

### Progressive ratio testing for cocaine IVSA and FSA

Following 10 days of FR1 testing, the mice in the cocaine and FSA studies were tested under a progressive ratio schedule of reinforcement in one session. The first reinforcer required one press of the active lever, and the response requirement doubled every reinforcer thereafter. The session ended when the mouse failed to earn a reinforcer in 30 minutes, or 2 hours total elapsed. The last ratio achieved was determined for each mouse and utilized as an indicator of performance.

### Acute locomotor effects of cocaine

Mice, which were naïve to any prior experimentation, were tested in an acute locomotor dose-response procedure. Testing occurred in Med Associates open field boxes (43.2X43.2X30.5 cm, Med-Associates MED-OFAS-RSU; St Albans VT6), housed individually in sound-attenuating chambers. All testing sessions were preceded by a 30-minute acclimation period in the open field box with no prior injection. Following this 30-minute session, mice were briefly removed and received an intraperitoneal (ip) injection of either saline (sessions 1 and 2) or cocaine (sessions 3, 4, and 5) at a dose volume of 10 mL/kg body-weight, and returned to the open field box for 1 hour. Cocaine was administered at 3 doses (5, 10, and 20 mg/kg body weight) in a within-subjects dose-response design. All 6 possible dose orders were utilized and balanced across genotype groups. Sessions 1, 2, and 3 occurred over 3 consecutive days. Sessions 4 occurred 2 days following session 3 and session 5 occurred 3 days following session 4, in order to limit any potential effects of prior cocaine exposures.

Locomotor behavior was assessed by distance traveled, as determined by infrared beam breaks. Distance traveled under no injection, saline injection, and cocaine injection was assessed. The acute locomotor effect of cocaine was calculated by subtracting the average of the distance traveled after both saline injection sessions from the distance traveled after cocaine injection. The data were binned into two, 30-minute bins and assessed within bin in addition to the full 1-hour session.

### Pre-experiment habituation

For all mice used in the behavioral tests described below, daily handling and habituation was initiated a week before the behavioral testing began. Mice were picked up by hand, stroked and touched for approximately 2 min per session. A quick assessment was made during the last handling session; if a mouse exhibited high levels of anxiety-like behavior (incontinence, hyperactivity, etc.), 1-2 additional handling sessions were conducted before experimental testing.

### Open Field Test

The open field test was conducted as described (*11*) using a 40 x 40 x 40 cm (l x w x h) square arena with white plastic boards. Each mouse was habituated within the procedure room for 30 minutes and was then placed in the center of the open field arena. The total distance travelled and duration of time in the center (∼25% of total area) or edge over a period of 15 minutes were recorded and analyzed using Viewer III software (BIOBSERVE, Bonn, Germany). The total duration of the open field test was 15 min.

### Novel Object Recognition Test

The novel object recognition test was conducted as described (*12*) with the same apparatus that used for the open field tests. A test mouse was given free access to the entire chamber for a 5 min habituation period. When the training session started, two identical objects (Lego blocks or a flask filled with bedding) were placed in diagonally opposite corners of the arena (6 cm from the wall), and the test mouse was allowed to freely explore for 10 min. After 24 hr, the test mouse was returned to the center of the arena and habituated for 5 min. When the testing session started, one familiar and one novel object were presented at the same positions in the arena. The test mouse was then given 10 min to explore the objects, while exploratory behaviors (sniffing, rearing against the objects, and head within 2 cm of the object) were recorded. Videos were processed using Viewer III tracking software by experienced personnel. The first 5 minutes of the training and testing sessions were used for analysis. The discrimination index (DI) was calculated as (Exploration time with novel object - Exploration time with familiar object) / (Exploration time with novel object + Exploration time with familiar object).

### Elevated Plus Maze Test

The elevated plus maze test was conducted as described (*11*) in an arena with two open arms and two closed arms that are raised 50 cm above the floor. All arms were 30 x 5 cm (l x w) with white walls (15 cm height) and floors. The test started 2 hr after the end of the dark cycle. Thirty minutes after acclimation in the behavioral room, the test mouse was placed in the center of the maze and allowed to freely explore the arena. The total test duration was 5 min. The duration of time spent in exploratory behaviors, the number of entries and the distance travelled in the open and closed arms (excluding center) were analyzed using Viewer III tracking software.

### Rotarod Test

Motor coordination was evaluated using the methods that are described in the Standard Operating Procedures of the Jackson Laboratory Mouse Neurobehavioral Phenotyping Facility using a five-station rotarod treadmill (ENV-575M, Med Associates, St. Albans, VT). Mice were first acclimated to the behavioral room for 1 hr before testing. The testing session consisted of three trials; in each trial, the speed was increased from 4 to 40 rpm; and each trial was separated by an interval of 1 min. A trial was terminated when a mouse fell off, clung to the rod and completed full passive rotation, or after 300 sec. The duration and end speed on rotarod were recorded and averaged from the last two trials for each mouse.

### Barnes Maze Test

Barnes maze tests were conducted as previously described (*13*). The experimental protocol consisted of a habituation session (day 1); 12 training sessions with 3 trials per day and a 15 min intertrial interval on days 2-5; and the testing session on day 6. During the training sessions, the mice were released into the middle of the maze, and they learned to enter the open escape hole to avoid exposure to a strong light. Three visual cues were placed at distinct positions outside of the maze to facilitate learning. The maze was cleaned with 70% ethanol thoroughly between trials to eliminate olfactory cues. Twenty-four hours after the training sessions, mice were tested in the arena for 90 sec with all holes closed. The result evaluated include primary errors (errors made before reaching the escape hole), latency (the time elapsed before reaching the escape hole), track length (the total length traveled) and target hole preference (percentage of time spent adjacent to the escape hole). Their performance was recorded and analyzed using Viewer III tracking software.

### The morphine Conditioned Place Preference (mCPP) Test

The mCPP test is performed according to previously described methods (*14*) using an activity-measuring system and two compartments that are separated by a clear plastic divider. One compartment has circular patterns on the walls and a smooth plastic black floor, whereas the other compartment has walls with stripped patterns and a white dotted plastic floor. On day 0, the mice are habituated in the measuring system for 20 min with free access to both compartments through a round hole in the separator. The duration and activities in each compartment are recorded. Mice are then randomly assigned to either the black or white compartment for morphine administration on the following days. On days 1, 3 and 5, mice received 0.1 ml intraperitoneal injections of 0.9% sterile saline; and are then placed into the assigned saline-compartment for 1 hr. Access to the other compartment was blocked by a clear plastic separator. On days 2, 4 and 6, the mice are placed in the alternative compartment for 1 hr immediately after receiving an intraperitoneal injections of 5 mg/kg morphine sulfate. On day 7, the mice are granted free access to both compartments for 20 min without receiving an injection. The duration and their activities in each compartment are recorded and analyzed using Activity Monitor software (Med Associates, Inc., St. Albans, VT).

### Opioid-induced Hyperalgesia

Opioid-induced hyperalgesia was evaluated using von Frey filaments (Product #58011, Soelting Co., Wood Dale, IL, USA) and the ‘up-down’ algorithm as previously described (*15, 16*). For these measurements, mice were individually habituated within a clear plastic cylinder placed upon a wire mesh platform for 30 min before a measurement was made. The cylinders were covered with black tile to prevent jumping and to reduce light exposure. After 30 min of habituation, the testing was initiated with a 0.6 g von Frey filament. When the test mouse was not walking or rearing, the filament tip was pressed against the mid-plantar surface of one hind paw to cause a slight bend in the filament and was left in place for 3 seconds. Withdrawal responses (flicking, withdrawal or licking of the stimulated paw) or no responses were recorded on a sheet provided by the Up-Down Reader. If a response was observed, then a filament of a slightly less weight was applied to the same paw; while a filament of larger weight was applied if no response observed. The collected data was then processed with open-source UDReader Software (*16*).

### MRI analyses

The brains of age-matched adult male C57BL/6 and *Nav1* KO mice (n=4/group) were examined by *in vivo* MRI using a high-field 7T MRI scanner (Bruker, Billerica, MA), which is housed at the Stanford Center for Innovation in In vivo Imaging (SCi^3^) facility. All mice were anesthetized under 1.0% to 1. 5% isoflurane that was administered by nose cone throughout the session. Their body temperatures were supported with warm air, while their respiratory rates were continuously monitored. Anatomical images were acquired using T2-weighted turbo rapid acquisition with relaxation enhancement (T2 TurboRARE) with the following parameters: repetition time (TR) = 2500 ms, echo time (TE) = 40 ms, flip angle = 90 degrees, slice thickness = 0.5 mm. Slices were obtained in the axial view (coronal in the mouse) with the first slice starting at the rostral-most extension of the prefrontal/motor cortex, while the olfactory bulb was excluded. The DICOM files obtained were processed using Osirix software (Pixmeo SARL, Bernex, Switzerland). The cortical thickness, hippocampal volume and brain volume were manually labelled by an experimenter that was blinded to the genotype of mice. Measurements were obtained from a continuous series of slices (for cortical thickness: from +0.63 to -0.67mm; for hippocampal volume: -0.77 to -3.37mm; for brain volume: +3.33 to -4.87mm; locations relative to Bregma) that were aligned across mice groups. The normalized hippocampal volume was calculated by dividing the absolute hippocampal volume by the total brain volume. The data were analyzed using Prism 9.1.0 (GraphPad Software, Inc. La Jolla, CA) with unpaired student t-test.

### Preparation of brain slices

Brain slices were prepared from anesthetized mice using previously described techniques (*17*). In brief, coronal slices (∼300 μm) were prepared from excised brains that were sectioned with a vibratome in cold (4 °C) “slicing” buffer (ACSF) containing: 126 mM NaCl, 2.5 mM KCl, 1.25 mM NaH_2_PO_4_, 1 mM CaCl_2_, 2 mM MgSO_4_, 26 mM NaHCO_3_, and 10 mM glucose; pH 7.4, when saturated with 95% O_2_/5% CO_2_. Slices were then transferred to an incubation chamber filled with standard ACSF buffer containing: 126 mM NaCl, 2.5 mM KCl, 1.25 mM NaH_2_PO_4_, 2 mM CaCl_2_, 1 mM MgSO_4_, 26 mM NaHCO_3_, and 10 mM glucose. The slices were incubated at 33 ± 1°C for 1 hr, and then at room temperature before use.

### Whole cell patch-clamp recording

After incubation, slices were transferred to a recording chamber where they were minimally submerged (32 ± 1°C) and perfused at the rate of 2.5-3 mL/min with standard ACSF buffer. Patch electrodes pulled from borosilicate glass tubing (1.5 mm OD) and had impedances of 4-6 MΩ when filled with Cs-gluconate based intracellular solution containing: 120 mM Cs-gluconate, 10 mM KCl, 11 mM EGTA, 1 mM CaCl_2_, 2 mM MgCl_2_, 10 mM HEPES, 2 mM Na2ATP, 0.5 mM NaGTP. The osmolarity of the pipette solution_−_ was adjusted to 285–295 mOsm and the pH to 7.35-7.4 with CsOH and ECl was -70 mV calculated from the Nernst equation. Whole cell voltage clamp recordings of miniature (m) IPSCs were obtained from the granule cells in the dentate gyrus of right hippocampus at a holding potential (V_h_) of +20 mV in the presence of 1 μM tetrodotoxin (TTX, Ascent Scientific) without application of glutamate receptor blocking agents (*17*). Miniature (m) EPSCs were recorded from the granule cells at V_h_ = −70 mV, the estimated E_Cl_^−^with the Cs-gluconate internal solution (*17*). All recordings were made with a Multiclamp 700A amplifier, sampled at 10 kHz, filtered at 4 kHz with a Digidata 1320A digitizer, and analyzed using Clampfit 9.0 (Molecular Devices, Sunnyvalle, CA), Mini Analysis (Synaptosoft, Decatur, GA), and Prism (GraphPad software). Only recordings with a stable access resistance <20 MΩ that varied <15% during the recording were accepted for analysis. One or two neurons were recorded per slice, and no more than three slices were used per mouse.

### Preparation of nuclei for snRNA-Seq analysis

This protocol is modified from the one developed by 10X Genomics. Brain tissue was obtained from age-matched adult male *Nav1* KO (n=4) and C57BL/6 (n=5) mice. The PFC was quickly dissected from the freshly extracted brain tissue and it was place in chilled Hibernate AB Complete (**HEB**) medium (BrainBits LLC, Springfield, IL) at 4 ^0^C. Freshly dissected PFC from each group were pooled in separate 50 ml conical tubes with a minimum amount of chilled Hibernate AB Complete (**HEB**) medium (BrainBits LLC, Springfield, IL). Then, 5 ml of chilled lysis buffer (2mM Tris-HCl, 2mM NaCl, 0.6 mM MgCl_2_ and 0.02% Nonidet^TM^ P40 Substitute in nuclease-free water) was added to the tubes, which were then incubated at 4 ^0^C for 10 min. The amount of lysis buffer and the incubation time were optimized determining the amount of buffer and incubation time that produced high-quality nuclei. After adding 5 more ml of HEB medium to the tubes, the tissues were triturated with a fire-polished silanized Pasteur pipette for 10-15 passes and then strained with 30 um MACS strainer (Miltenyi Biotec Inc, Auburn CA). Nuclei were pelleted by centrifugation at 500xg for 5 min at 4°C and were then resuspended in a chilled PBS with 1% BSA (Invitrogen AM2618, Thermo Fisher Scientific, Pittsburgh PA) wash buffer with 0.2 U/ul RNase inhibitor (#3335402001, Millipore Sigma, Darmstadt Germany). The nuclei were pelleted and resuspended twice, strained using a 30 um MACS strainer, and then centrifuged and resuspended in a chilled wash buffer. Mouse anti-NeuN antibody (MAB377, Millipore Sigma, Darmstadt Germany) was added at a 1:500 dilution, and the samples were incubated for 40 min at 4°C. Samples were then centrifuged at 400xg for 5 minutes at 4°C and the pelleted nuclei were resuspended with chilled wash buffer. Alexa 647 chicken anti-mouse antibody (A21463, Life technologies, Pittsburgh PA) was added to the preparations at a 1:500 dilution and the samples were incubated for 40 min at 4°C. After incubation, nuclei were pelleted and resuspended, and Hoechst 33342 (Invitrogen H3570, Thermo Fisher Scientific, Pittsburgh PA) was added to a final concentration of 0.25 ug/ml.

Single nuclei sorting was conducted using a 6-laser BD Influx sorter (BD Biosciences, San Jose CA) with a 100-um nozzle in the Stanford Shared FACS Facility. Various control samples (unstained, Hochest33342^+^ or NeuN^+^) were examined to optimize the gating strategy. Nuclei were gated based on size, scatter properties and staining for Hoechst and NeuN. To ensure that we were able to analyze different types of cells, ∼60% of the sorted nuclei were collected as NeuN^+^ and 40% were NeuN^-^. Single nuclei were sorted into collecting tubes and then visually inspected under a microscope for quality control; they were then pelleted and resuspended to produce a solution with 600 nuclei/ul in the final volume. Single nuclei were then captured in droplets with barcodes using the 10x Genomics Chromium system and cDNA libraries were produced using Chromium Next GEM Single Cell 3’ Reagent Kits v3.1 (10x Genomics, Pleasanton, CA) according to the manufacturer’s instructions. The *Nav1* KO and C57BL/6 samples were sequenced on the NovaSeq platform (Illumina, San Diego CA).

### snRNA-Seq data analysis

FASTQ files with the snRNA-Seq data were processed using Cell Ranger software (v6.0.0) and the “*cellranger count*” pipeline to generate a filtered feature-barcode matrix (the gene expression matrix). The reads within the C57BL/6 and *Nav1* KO FASTQ files were aligned using the Cell Ranger built-in mouse reference (mm10). Two gene expression matrices were produced: 23445 features × 15402 cells for C57BL/6; and 23737 features × 14623 cells for *Nav1* KO. The C57BL/6 and *Nav1* KO gene expression matrices were then analyzed using the R/Seurat (v4.0.1) package. Low-quality cells with unique gene counts < 200 and >10% mitochondrial counts were filtered out. The high-quality C57BL/6 (20955 features × 14630 cells) and Nav1 KO (21310 features × 14056 cells) matrices were merged into one Seurat object (21892 features × 28686 cells) that was used for subsequent analyses (data normalization, variable feature identification, dimensional reduction, etc.).

The gene counts for each cell within the global matrix was normalized by dividing it by the total counts in that nucleus; and this number was multiplied by 10000 and were then natural log transformed to become the normalized values. We also identified 2000 variable features, which exhibited high cell to cell variability in the matrix. The matrix was then further scaled for linear dimensional reduction purposes. For the scaled gene expression matrix, the mean expression of a gene across different cells is set to 0 and the variance is set to 1. Principal component analysis (PCA) was performed on the scaled data, and the first 10 PCs were chosen to represent the dimensionality of the global matrix. The default K-nearest neighbor (KNN) graph-based method and Louvain algorithm for single-cell clustering was used; and the resolution parameter was set to 0.3. In total, 14 clusters (cluster 0 to 13) were classified; and of these, cluster 0 contained the most cells while cluster 13 contained the fewest. To visualize the cells in low-dimensional space (to aid interpretation), the non-linear dimensional reduction program (UMAP) (*18*) was performed using the first 10 PCs.

The differentially expressed genes (**DEG**) for each cluster were detected using a minimum percentage of 0.25 in either of the two clusters and an average log2 fold-change (FC) ≥ 0.25. Cell type specific canonical markers were used to determine the cell type identity of the 14 clusters. Non-neuron cells were assigned as follows: Astrocyte (*Gja1, Aqp4*), Microglia (*C1qa*), and Oligodendrocyte (*Apsa, Mbp*). Of note, no endothelial cell markers (*Flt1, Cldn5, Nostrin*) were highly expressed in any of 14 clusters. The type of neuronal cells was determined by whether they expressed mRNA markers for excitatory (*Slc17a7, Tshz2, Thsd7a*) or inhibitory (*Gad1, Gad2, Slc32a1, Meis2*) neuronal markers. To verify the cell type assignments, our C57BL/6 gene expression matrix was compared with that of a published reference (GSE124952) (*19*) data set for PFC cells obtained from saline control C57BL/6 mice, which generated an expression matrix with 20718 features × 11886 cells. The canonical correlation analysis (CCA) between this reference and our C57BL/6 dataset was used to remove batch effects before matrix integration. We then projected the reference single cells with their cell type labels onto the UMAP plot for comparison with our C57BL/6 dataset.

### Tissue collection and sectioning

After isoflurane anesthesia, transcardial perfusion (Harvard Apparatus p-70, Holliston, MA) was performed with a 0.15 M NaCl solution was followed by fixation in 4% paraformaldehyde (Aldrich Chemistry, Darmstadt, Germany). The brains were then extracted and post-fixed in 4% paraformaldehyde for 4 h. After fixation, the brains were transferred into a 30% sucrose solution, and then stored at 4°C until sectioning. The brain tissue slices used for immunohistochemistry were sectioned at 30 µm, and then stored in a cryo-protective solution (PBS, 20g PVP-40, 600ml ethylene glycol, 600g sucrose) at -20°C.

### Immunohistochemistry

The immunohistochemical analyses were performed as described in (*20*). In brief, brain sections were rinsed 5 times in PBS (Sigma-Aldrich P5368-10pak) for 5 min, and then blocked in 10% normal donkey serum and 0.3% Triton X-100 in PBS to minimize nonspecific binding. For c-Fos staining, the brain sections were heated to 80 °C in 50 mM Citrate buffer (PH 6.0) for 20 minutes to expose epitopes for antibody binding. The sections were then incubated with the primary antibodies (**Table S3**) in the blocking solution for 48 hr at 4°C. The sections were then rinsed in PBS and incubated with a corresponding secondary antibody for 4 hr. After the last rinse, the sections were mounted, air dried overnight; and were then sealed with a cover slip in Dako Fluorescence Mounting Medium (S3023, Dako North America, Inc., Carpinteria, CA) and stored at -20°C.

### Image Capture

Images were captured and analyzed using a Leica SP8 white light pulsed laser confocal microscope and Leica LAS X Premium software at the Cell Sciences Imaging Facility at Stanford. Channels were selected and the exposure times were adjusted to optimize the images. Imaris software (Bitplane Inc., Concord, MA) was used for image capture (*21*).

### Brain slice preparation

Mice were anesthetized, and using previously described techniques (*17*), brains were removed, and coronal slices (∼300 μm) were cut with a vibratome in cold (4 ± 1°C) “slicing” ACSF containing (in mM): 126 NaCl, 2.5 KCl, 1.25 NaH_2_PO_4_, 1 CaCl_2_, 2MgSO_4_, 26 NaHCO_3_, and 10 glucose; pH 7.4, when saturated with 95% O_2_/5% CO_2_. Slices were then transferred to an incubation chamber filled with standard ACSF containing (in mM): 126 NaCl, 2.5 KCl, 1.25 NaH_2_PO_4_, 2 CaCl_2_, 1 MgSO_4_, 26 NaHCO_3_, and 10 glucose. Slices were incubated at 33 ± 1°C for 1 h and then at room temperature before use.

### Whole cell patch-clamp recording

After incubation, slices were transferred to a recording chamber where they were minimally submerged (32 ± 1°C) and perfused at the rate of 2.5-3 mL/min with standard ACSF. Patch electrodes pulled from borosilicate glass tubing (1.5 mm OD) and had impedances of 4-6 MΩ when filled with Cs-gluconate based intracellular solution containing (in mM): 120 Cs-gluconate, 10 KCl, 11 EGTA, 1 CaCl_2_, 2 MgCl_2_, 10 HEPES, 2Na2ATP, 0.5 NaGTP. The osmolarity of the pipette solution was adjusted to 285–295 mOs_−_ and the pH to 7.35-7.4 with CsOH and ECl was -70 mV calculated from the Nernst equation.

Whole cell voltage-clamp recordings of miniature (m) IPSCs were obtained from the granule cells in the dentate gyrus of right hippocampus at a holding potential (V_h_) of +20 mV in the presence of 1 μM tetrodotoxin (TTX, Ascent Scientific) without application of glutamate receptor blockers (*17*). Miniature (m) EPSCs were recorded from the granule cells at V_h_ = −70 mV, the _−_ estimated EC with the Cs-gluconate internal solution (*17*). All recordings were made with a Multiclamp 700A amplifier, sampled at 10 kHz, filtered at 4 kHz with a Digidata 1320A digitizer, and analyzed using Clampfit 9.0 (Molecular Devices, Sunnyvalle, CA), Mini Analysis (Synaptosoft, Decatur, GA), and Prism (GraphPad software). Only recordings with a stable access resistance <20 MΩ that varied <15% during the recording were accepted for analysis. One or two neurons were recorded per slice, and no more than three slices were used per mouse.

### Statistics

Data were analyzed and graphed using Prism 8.4.1 for Windows (GraphPad Software, Inc. La Jolla, CA), Statistica 8.0 (TIBCO Software, Inc. Palo Alto, CA) or SPSS (IBM Corp. Released 2020. IBM SPSS Statistics for Windows, Version 27.0. Armonk, NY: IBM Corp). The cocaine IVSA, FSA and acute cocaine locomotor behavior results were assessed by ANOVA. Interactions and main effects were decomposed by simple main effects and pairwise comparisons with Sidak correction for multiple comparisons. Since cocaine IVSA data tends to depart from normal distributions, all cocaine IVSA data was log10 transformed prior to analysis. For the mCPP, acute locomotor response and hyperalgesia, Barnes maze test and novel object recognition test, the statistical significance was determined using a two-factor ANOVA with repeated measures. Stimulus (morphine vs. saline; familiar vs novel) or genotype (control vs. KO) were the two factors assessed. Post-hoc analysis was conducted using a Bonferroni post-test. Exploratory behaviors (including the EPM and open field tests) were evaluated using a one-way ANOVA with Tukey’s post-test. In the figures, the significance levels are indicated by: * p<0.05, ** p<0.01, *** p<0.001, **** p<0.0001.

---

**Table S1.** The results of association tests for population structure (PS) that were performed on haplotype blocks within the indicated genes are shown. The PS association test was performed using the method described in (22), and a p-value >0.05 indicates that the alleles within the haplotype block do not reflect population structure. The p-value for the genetic association with the cocaine IVSA data, which was calculated by the HBCGM program, is also shown for each haplotype block.

| **Gene** | **HBCGM p-value** | **PS p-value** |
| --- | --- | --- |
| *Tnni1* | $1.3 \times 10^{-06}$ | 0.16 |
| *Etnk2* | $1.8 \times 10^{-06}$ | 0.0032 |
| ***Nav1*** | $3.4 \times 10^{-06}$ | **0.26** |
| *Ipo9* | $4.2 \times 10^{-06}$ | 0.040 |
| *Sh3pxd2a* | $5.1 \times 10^{-06}$ | 0.11 |
| *Cntn2* | $8.5 \times 10^{-06}$ | 0.49 |
| *Chi3l1* | $1.2 \times 10^{-05}$ | 0.13 |

**Table S2.** Synonymous and nonsynonymous SNPs within the coding sequence of *Nav1*. The nucleotide position, the affected amino acid, and the C57BL/6 and the alternative (ALT) amino acids are shown for each SNP. The three cSNPs are highlighted in bold.

| <u>Chr</u> | <u>Genomic Pos</u> | <u>Amino Acid</u> | <u>C57BL/6</u> | <u>ALT</u> |
| --- | --- | --- | --- | --- |
| 1 | 135584853 | 156 | S | S |
| 1 | 135584727 | **198** | **D** | **E** |
| 1 | 135532666 | 306 | S | S |
| 1 | 135532552 | 344 | G | G |
| 1 | 135532354 | 410 | T | T |
| 1 | 135472430 | 467 | S | S |
| 1 | 135471052 | 597 | S | S |
| 1 | 135470854 | 663 | L | L |
| 1 | 135470761 | 694 | R | R |
| 1 | 135470033 | 799 | S | S |
| 1 | 135469724 | 902 | S | S |
| 1 | 135469698 | **911** | **A** | **V** |
| 1 | 135454498 | 1319 | S | S |
| 1 | 135454067 | **1366** | **P** | **L** |
| 1 | 135450851 | 1567 | D | D |
| 1 | 135450848 | 1568 | L | L |
| 1 | 135450076 | 1618 | S | S |
| 1 | 135450064 | 1622 | E | E |
| 1 | 135449968 | 1654 | L | L |
| 1 | 135449036 | 1683 | F | F |
| 1 | 135448967 | 1706 | I | I |
| 1 | 135448937 | 1716 | G | G |
| 1 | 135441768 | 1788 | L | L |

**Table S3.** The primary antibodies used for immunohistochemistry and their source and the dilution that was used in an experiment are shown.

| **Primary Antibody** | **Cat #** | **Vendor** | **Dilution** |
| --- | --- | --- | --- |
| guinea pig anti-vGlut1 | 135304 | Synaptic Systems | 1/1000 |
| rabbit anti-PSD95 | 51-6900 | Invitrogen | 7/1000 |
| guinea pig anti-vGat | 131004 | Synaptic Systems | 1/1000 |
| mouse anti-gephyrin | 147021 | Synaptic Systems | 1/400 |
| rabbit anti-MAP2 | AB5622 | millipore | 1/1000 |
| Mouse anti-tubulin | 801201 | BioLegend | 1/1000 |
| Rabbit anti-c-fos | Ab190289 | Abcam | 1/500 |

**Table S4.**
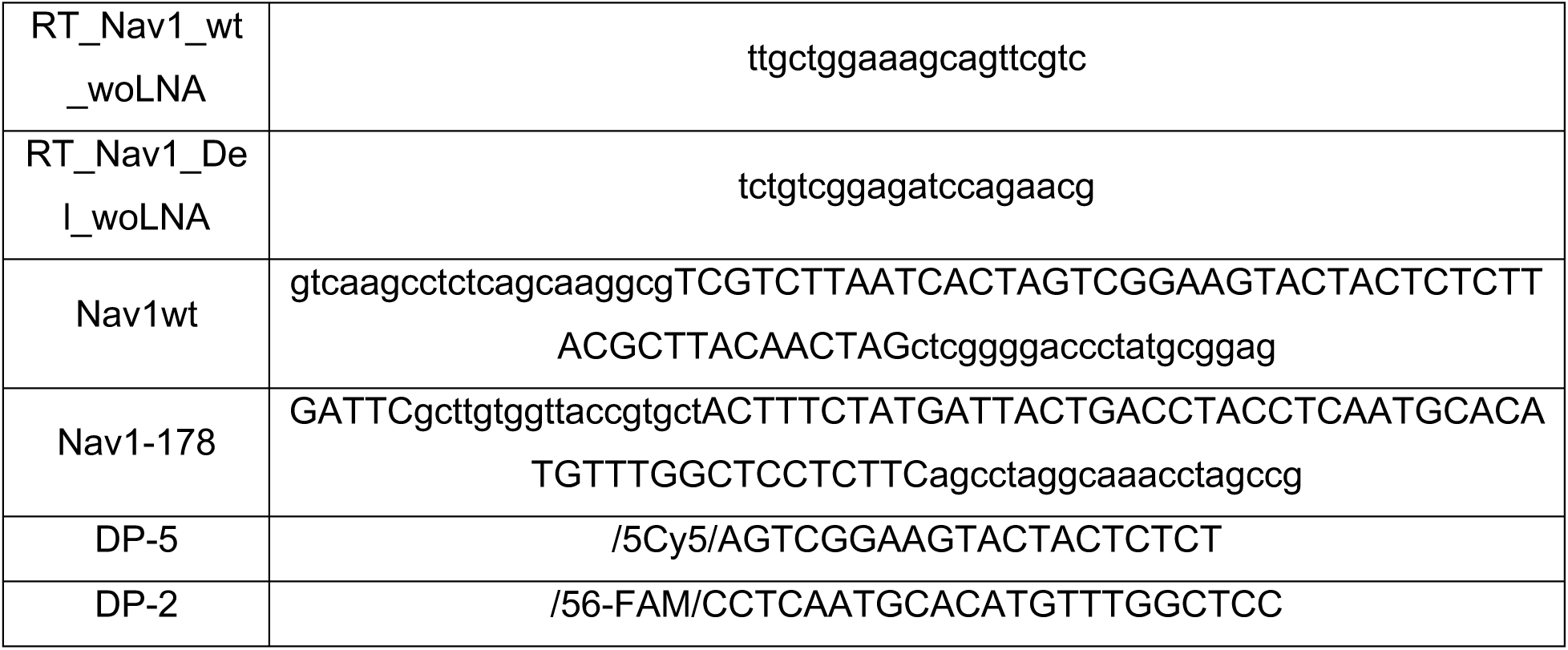
The oligonucleotide probes used for smFISH analysis are shown.

**Figure S1.**
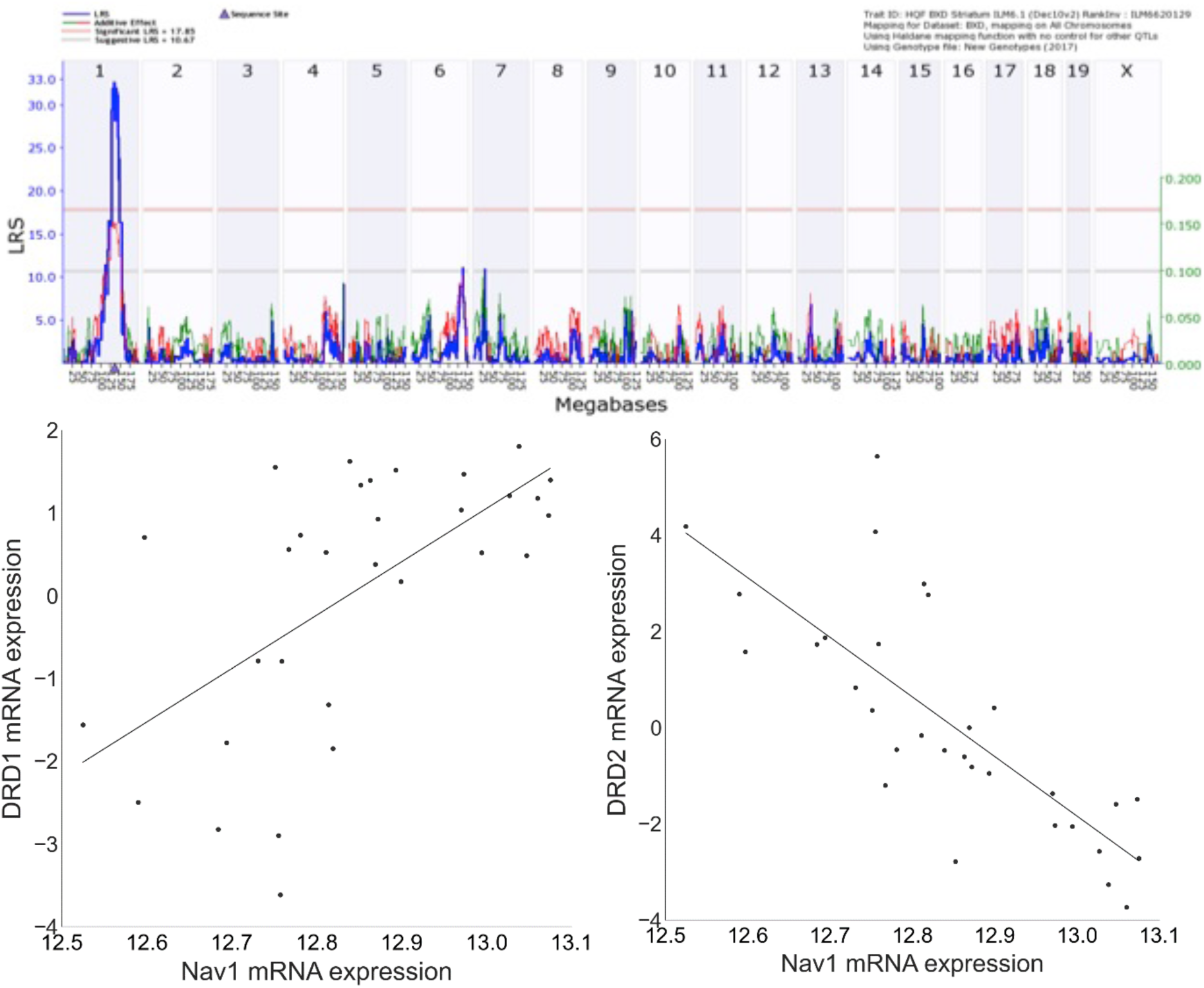
*Top:* The level of striatal *Nav1* mRNA expression in adult BXD mice (genenetwork.org; record ID ILM6620129) is controlled by cis-acting alleles within the *Nav1* locus (Chr 1 135.4 MB). The red horizontal line indicates the genome-wide significance threshold for an LRS/LOD score as determined by permutation analysis. *Bottom:* The *Nav1* eQTL was evaluated for genetically correlated phenotypes measured in the BXD database (genenetwork.org). The level of correlation of *Nav1* mRNA expression with the entire database of BXD phenotypes was evaluated. *Nav1* mRNA expression was most highly correlated (inverse) with the level of striatal *dopamine receptor D2 (Drd2*) mRNA expression (record ID: 15186; r=-.78; p=4.23x10^-8^). The *Drd2* mRNA correlation is quite specific because striatal *Drd1* mRNA expression is positively associated (record ID: 15186; r=.61; p=1.81x10^-4^) with the level of *Nav1* mRNA expression.

**Figure S2.**
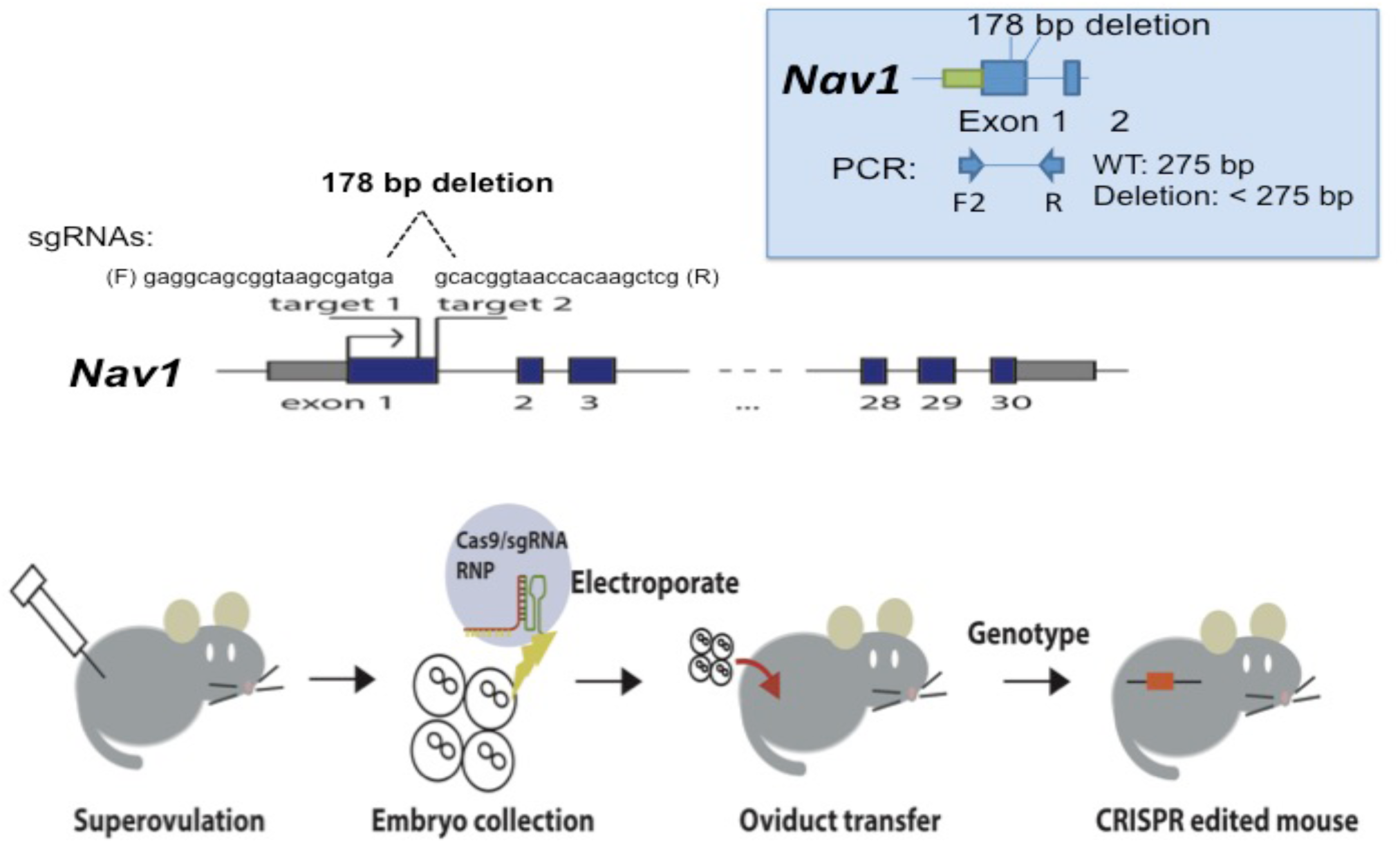
CRISPR-mediated genome engineering was used to produce *Nav1* knockout mice. C57BL/6 female mice were super-ovulated by intraperitoneal injection of pregnant mare serum gonadotropin and human chorionic gonadotropin. These mice were paired with C57BL/6 males to generate fertile embryos, and pronucleus stage embryos were collected. Then, Cas9, and the two sgRNAs (shown above) were electroporated into embryos. These sgRNAs were designed to delete a 178 bp region at the end of exon 1 in *Nav1*, and it also introduced an early stop codon into exon 1. Healthy embryos were then transferred into the oviducts of pseudo-pregnant recipient females. Genomic DNA from the resulting pups is screened by PCR amplification using the strategy shown in the colored box. While an intact *Nav1* gene will produce a 275 bp amplicon, genomic DNA from an engineered mouse will generate a shorter amplicon that is diagnostic of a deletion. Mice with a heterozygous *Nav1* KO were then bred to produce homozygous *Nav1* KO mice.

**Figure S3.**
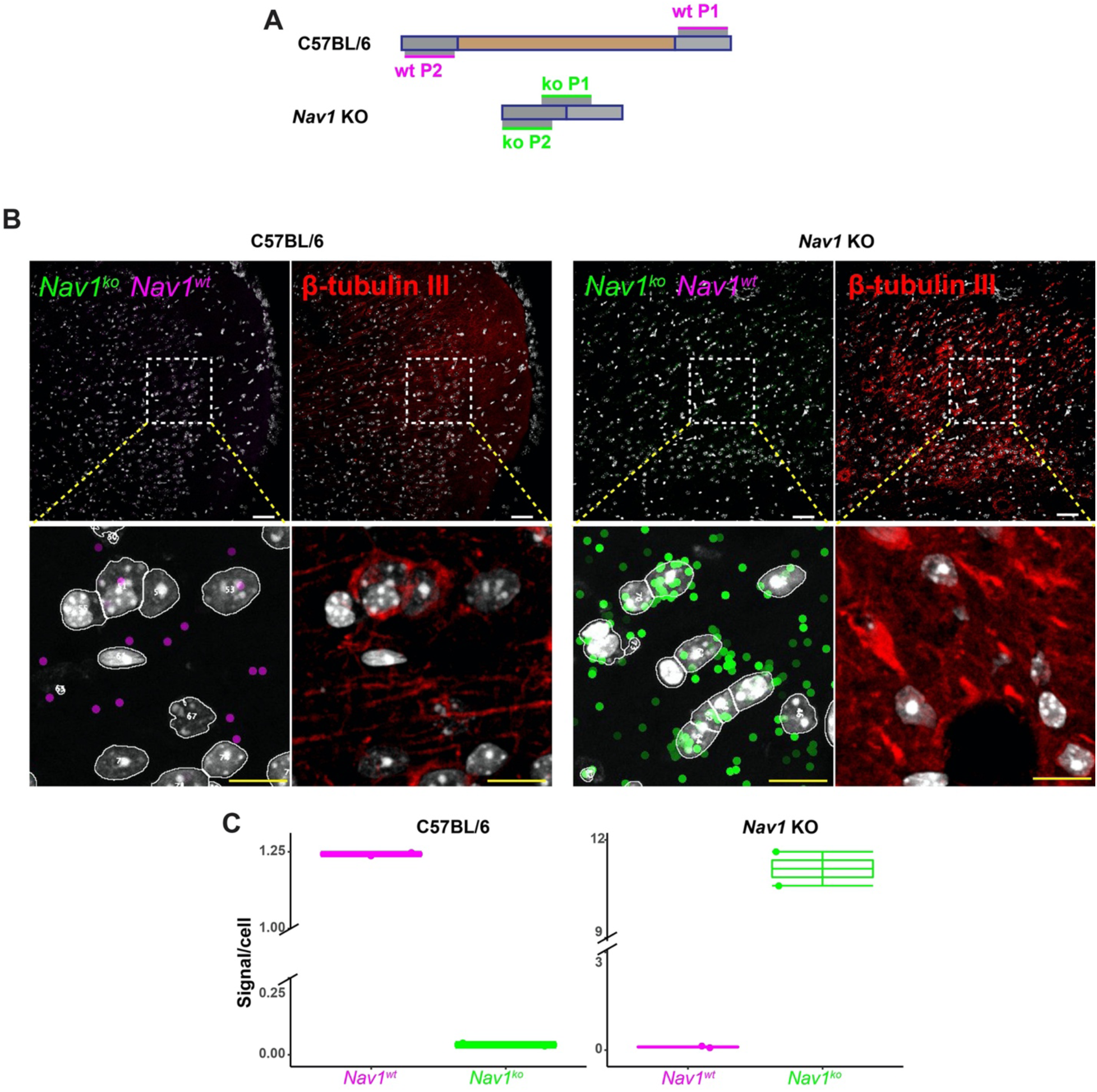
*Nav1* mRNA is not expressed in *Nav1* KO mice. **(A)** An illustration of the genomic regions within Exon 1 of *Nav1* recognized by the padlock probes that are specific for wildtype *Nav1* or *Nav1* KO mRNA (*9*). The gray boxes represent the 5’ and 3’ regions of exon 1 and the brown box represents the -178 bp deletion that is introduced into exon 1 to produce the *Nav1* KO. The wildtype *Nav1* probe (Nav1 wt) binds to sites (wtP1, wtP2) present in the wildtype *Nav1* allele; while the *Nav1* KO probe binds to adjacent regions around the 178 bp deletion (koP1, koP2), which are specific for the *Nav1* KO allele. **(B)** sm-FISH images obtained using the *Nav1* and *Nav1* KO probes on coronal sections of prefrontal cortex (PFC) tissue obtained from C57BL/6 and *Nav1* KO mice. The sections were also stained with anti-b-tubulin (b-Tubulin III) antibodies to identify neurons. The top panels show low power images; and the bottom panels are high power images of the boxed region shown in the top row. The scale bars are 50 µm in the upper images, and 10 um in the lower images. The staining pattern indicates that wild type *Nav1* mRNA is expressed in the PFC of C57BL/6 but not in *Nav1* KO mice, while the *Nav1* KO transcript is exclusively expressed in *Nav1* KO mice. **(C)** These boxplots show the single molecule counts generated for each padlock probe in the C57BL/6 and *Nav1* KO brain sections. Each group has 2 independent measurements.

**Figure S4.**
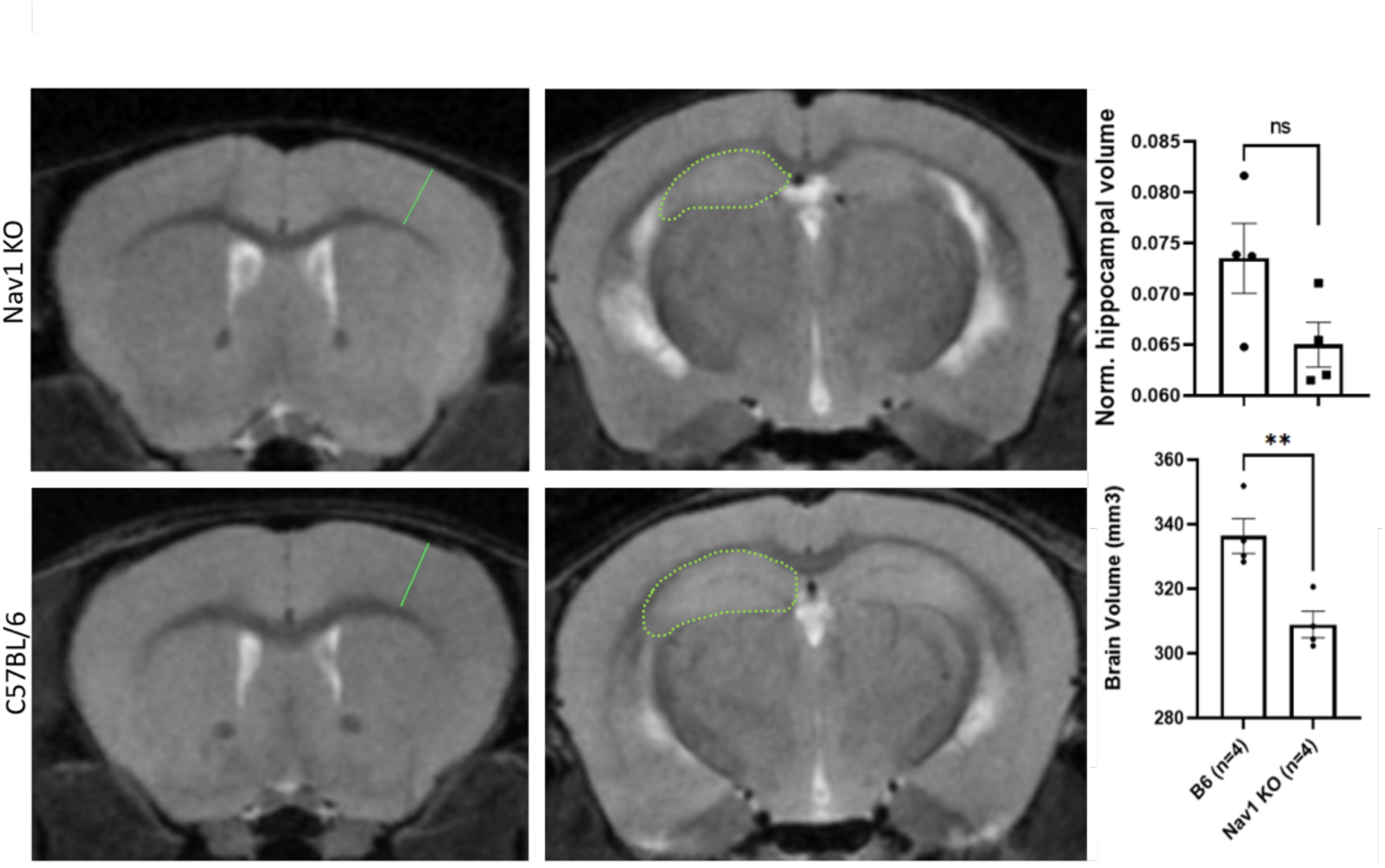
MRI scans of coronal sections of *Nav1* KO and age-matched isogenic C57BL/6 mice that were obtained with a Bruker 7-T MRI. The cortical thickness (solid line) and hippocampal volume (dotted line) measurements were generated as indicated in the images. In the adjacent graphs, each bar is the average ± SEM measurements made on *Nav1* KO and C57BL/6 (n=4 per group) mice. *Nav1* KO mice have a slightly smaller overall brain volume than C57BL/6 mice (p=0.007). However, after normalization relative to brain volume, there was not a significant difference (p=0.08) between the hippocampal volumes of C57BL/6 and *Nav1* KO mice.

**Figure S5.**
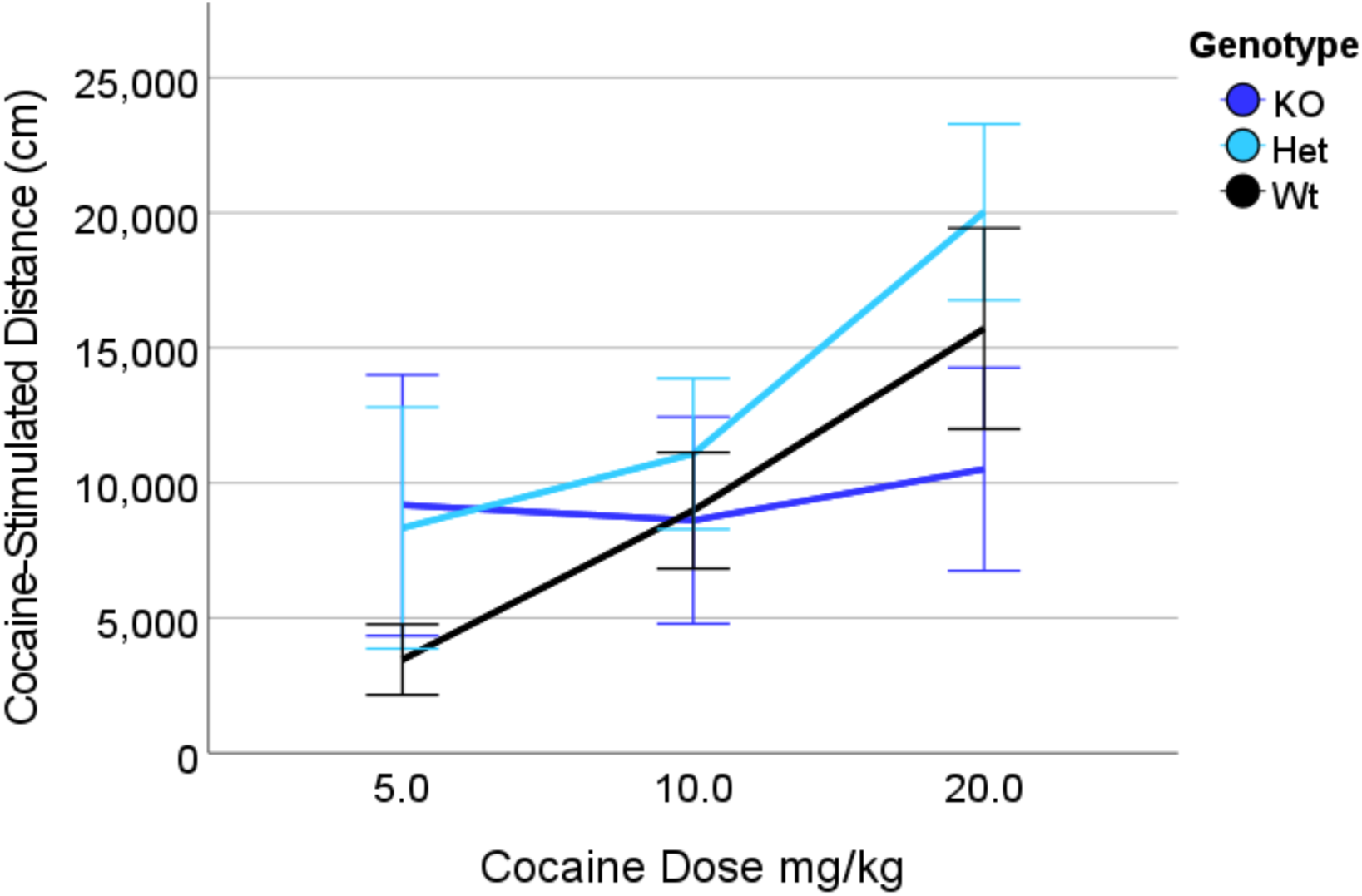
The acute locomotor response measured after cocaine injection exhibited by C57BL/6 (Wt), Het, and *Nav1* KO mice. The distance traveled over a 60 minute period after injection of each indicated dose of cocaine was increased (p<0.05). However, there were no significant differences in the extent of cocaine-induced locomotion among the mice with different genotypes (p>0.05). Each bar point is the average ±SEM of measurements made on 9-12 mice of each genotype.

**Figure S6.**
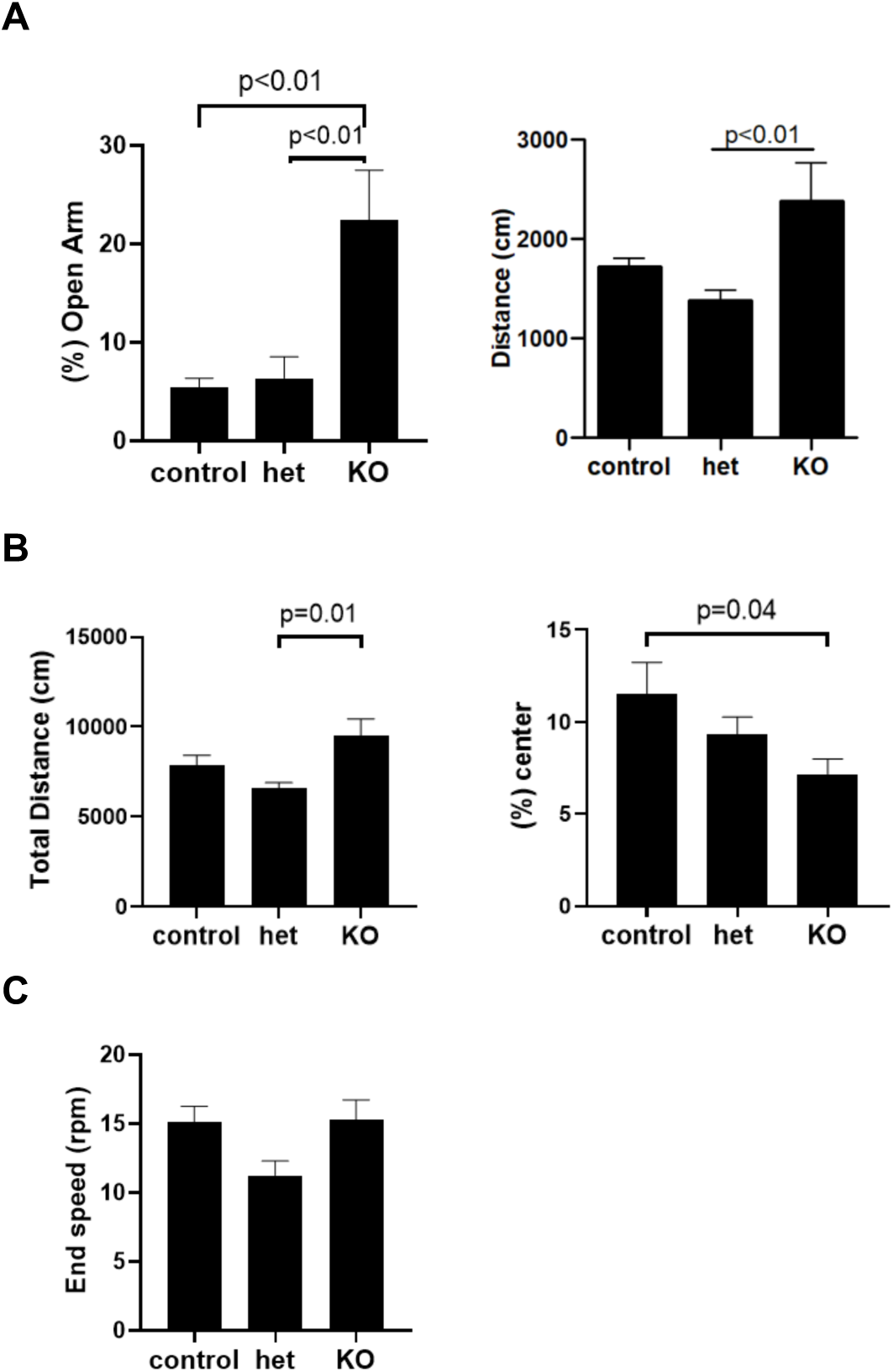
*Nav1* KO mice have increased basal locomotory activity, altered exploratory behavior, and normal motor coordination. (**A**) The exploratory **(left)** and locomotor activities **(right)** of C57BL/6 (control), het Nav1 KO and homozygous *Nav1* KO mice were measured in the elevated plus maze test. (**B)** In the open field test, *Nav1* KO mice travel a greater distance but spend less time in the center than control mice. (**C**) In the rotorod test, the motor coordination of Nav1 KO mice is like that of control mice (p>0.99). In these graphs, the bars are the mean ± SEM of measurements made on 5 – 7 mice per group. The p-values where there were significant differences between groups of mice are indicated above the graphs.

**Figure S7.**
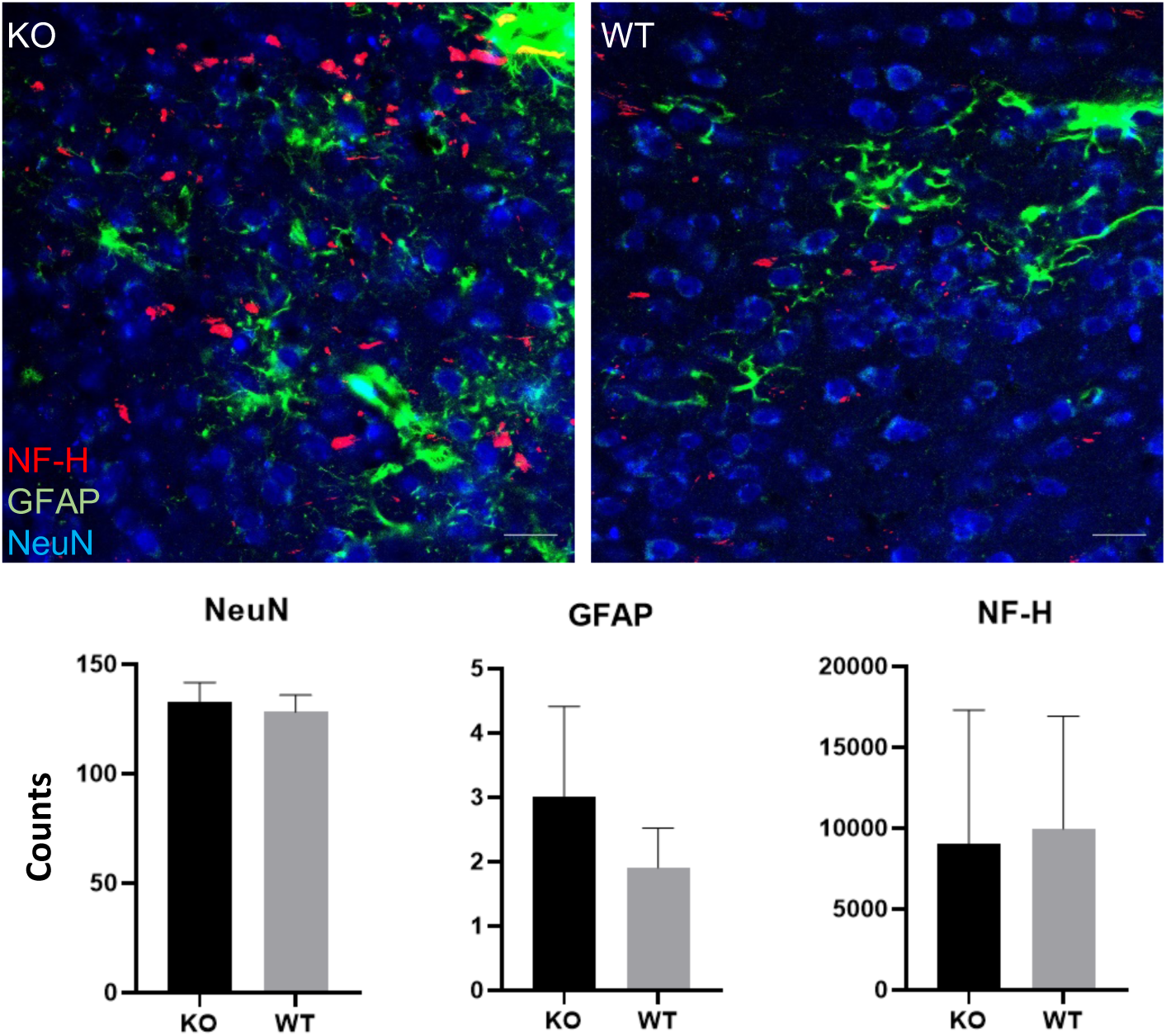
The cells in the nucleus accumbens were not altered *in Nav1* KO mice. *Top*: Confocal images of representative sections of the nucleus accumbens that were obtained from *Nav1* KO (left) and C57BL/6 (right) mice that were stained with antibodies to neurofilament heavy chain (NF-H), glial fibrillary acidic protein (GFAP), and neuronal nuclear protein (NeuN). Scale bar: 20 um. *Bottom*: Immunostained sections of the nucleus accumbens were analyzed using CellProfiler software (the Broad Institute of MIT and Harvard) (*23*). Each bar is the average ± SEM of analyses performed on 16 projection images (produced from 2.54 um z-stack) of 3 animals/group. There was no significant difference in the level of expression of NeuN (p value =0.53), GFAP (p value =0.28) or NF-H (p value =0.89) in the nucleus accumbens of *Nav1* KO (vs. C57BL/6 mice).

**Figure S8.**
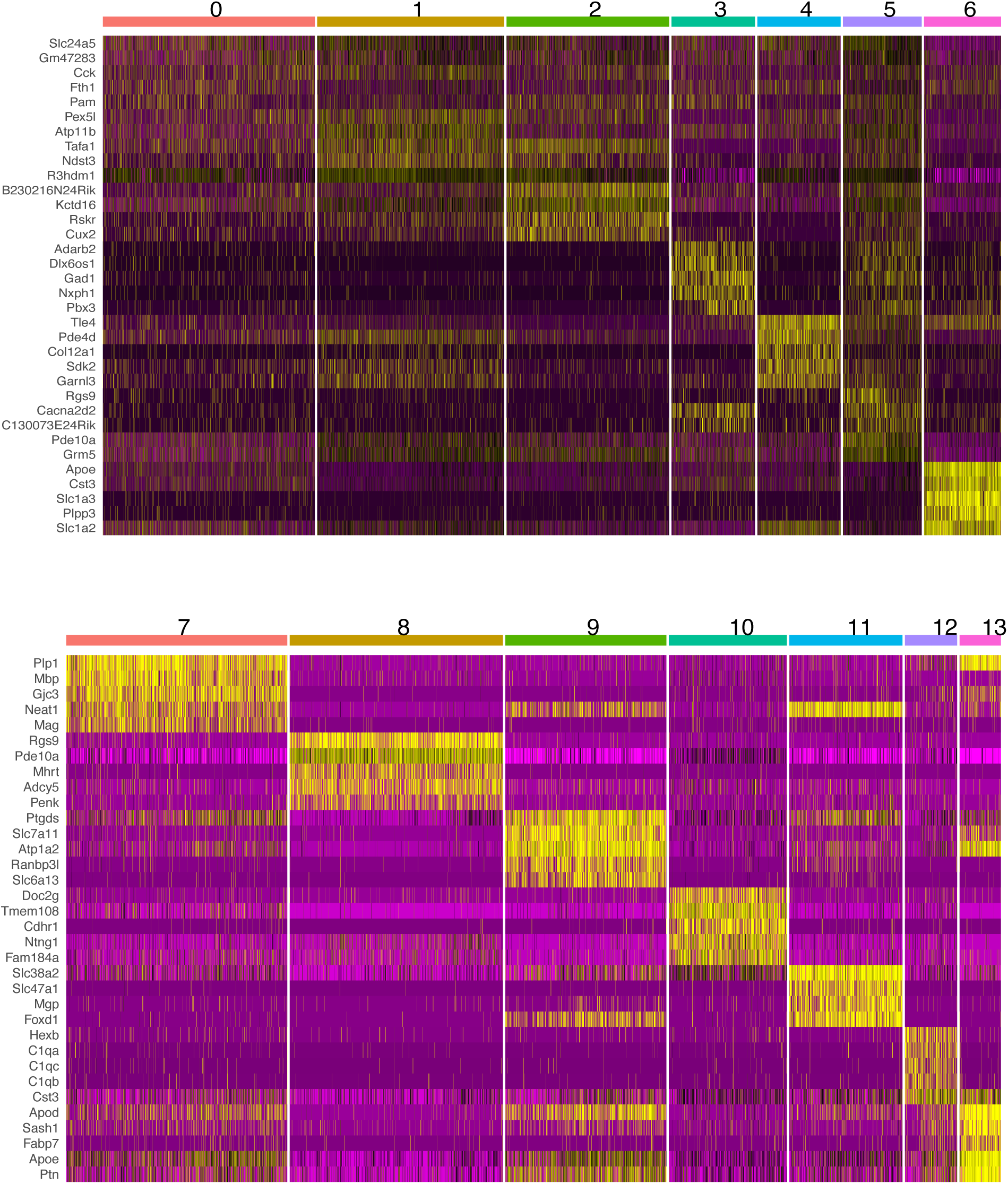
A heat map showing the 5 most differentially expressed genes (DEG) for each of the 14 clusters identified in the PFC samples analyzed. The gene symbols are shown on the left. The DEGs have a mean log2(fold-change) of 2.2 and a mean percentage of 0.65 in that cluster.

**Figure S9.**
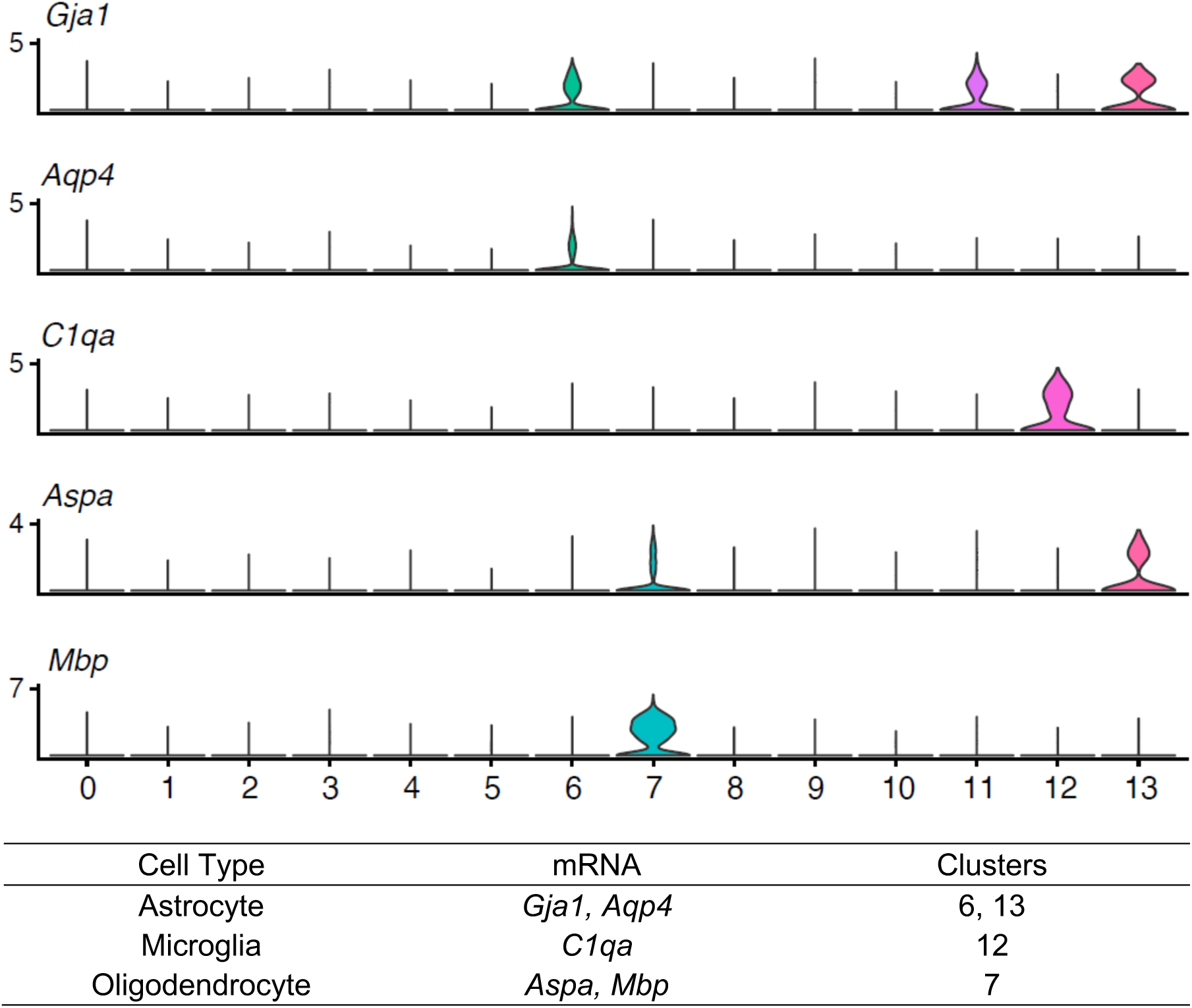
*Top*: Violin plots showing the level of expression of non-neuronal cell markers within each of the 14 cell clusters identified in the PFC. *Bottom:* The identity of the non-neuronal cell clusters was determined by their pattern of expression of the indicated canonical marker mRNAs (*19*). The cluster number is indicated on the x-axis and the y-axis shows the natural log transformed and normalized level of expression for each mRNA.

**Figure S10.**
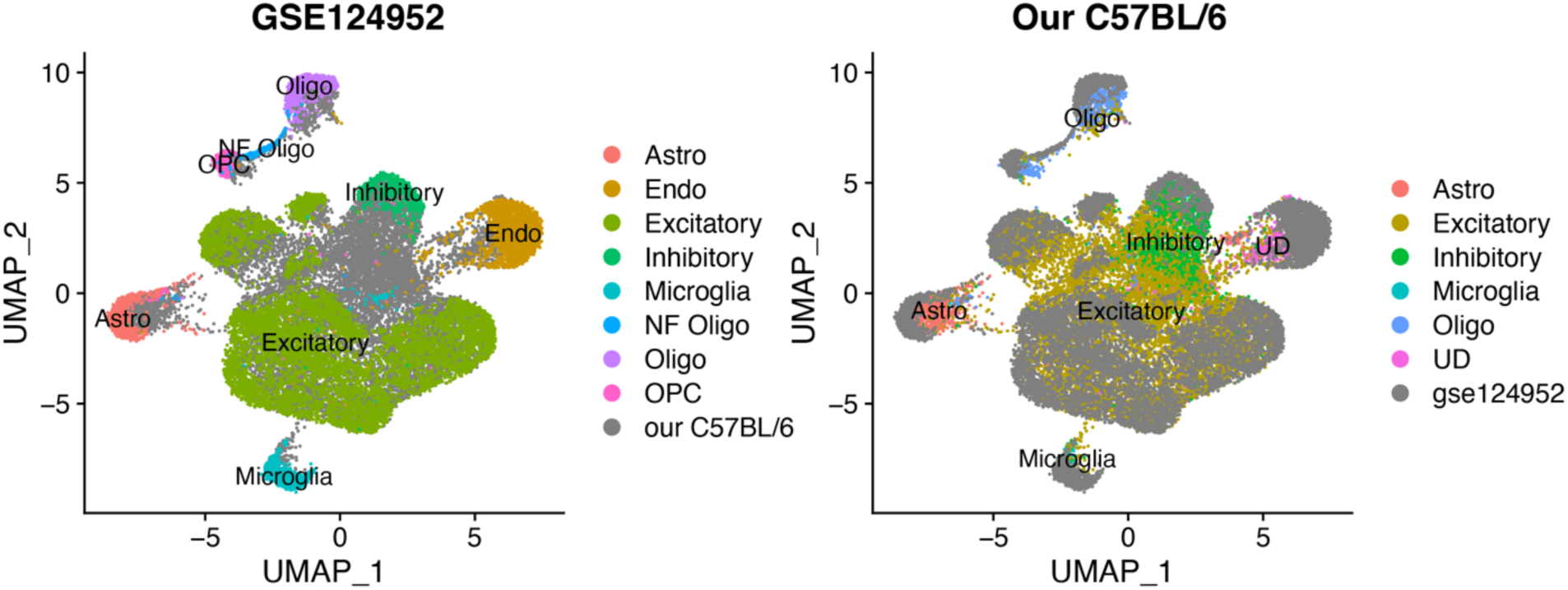
UMAP plots comparing a published scRNA-Seq dataset (GSE124952) (*19*) generated from PFC prepared from adult C57BL/6 mice with our C57BL/6 PFC snRNA-Seq data. Each plot presents the overlap of the cells from the two datasets. The cell types within the clusters for each dataset are indicated by the dot color. For comparison purposes, a gray dot on the left is a cell from our C57BL/6 dataset and a gray dot on the right is from a cell in the published dataset. These comparisons demonstrate the very high level of concordance between the cell clusters and their distribution in these two data sets.

**Figure S11.**
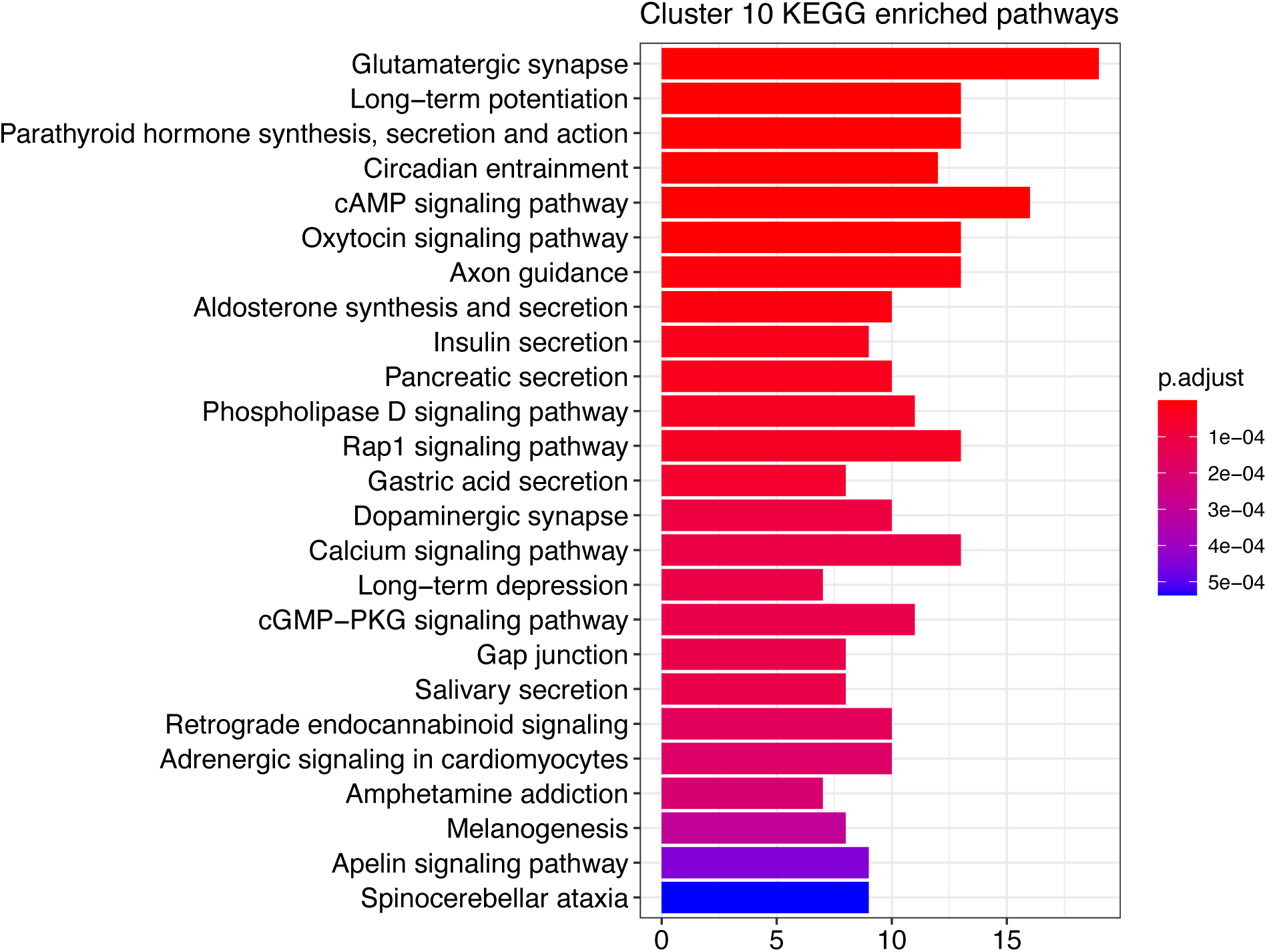
This barplot shows the 25 KEGG pathways that were most significantly enriched when the DEGs in cluster 10 were analyzed relative to those in other excitatory neuronal clusters (0-2, 4, 5). The bar length indicates the number of DEGs within the indicated pathway. The FDR controlled p-values, which were calculated using the Benjamini Hochberg method, are indicated by their color (as shown on the right). While many of the enriched pathways were associated with neuronal guidance, synapse and signaling functions; seven of the DEGs in cluster 10 (*Gria3, Grin2b, Grin2a, Camk2a, Ppp3ca, Gria4, Prkcb*) were associated with an amphetamine addiction pathway (p=0.000025).

**Figure S12.**
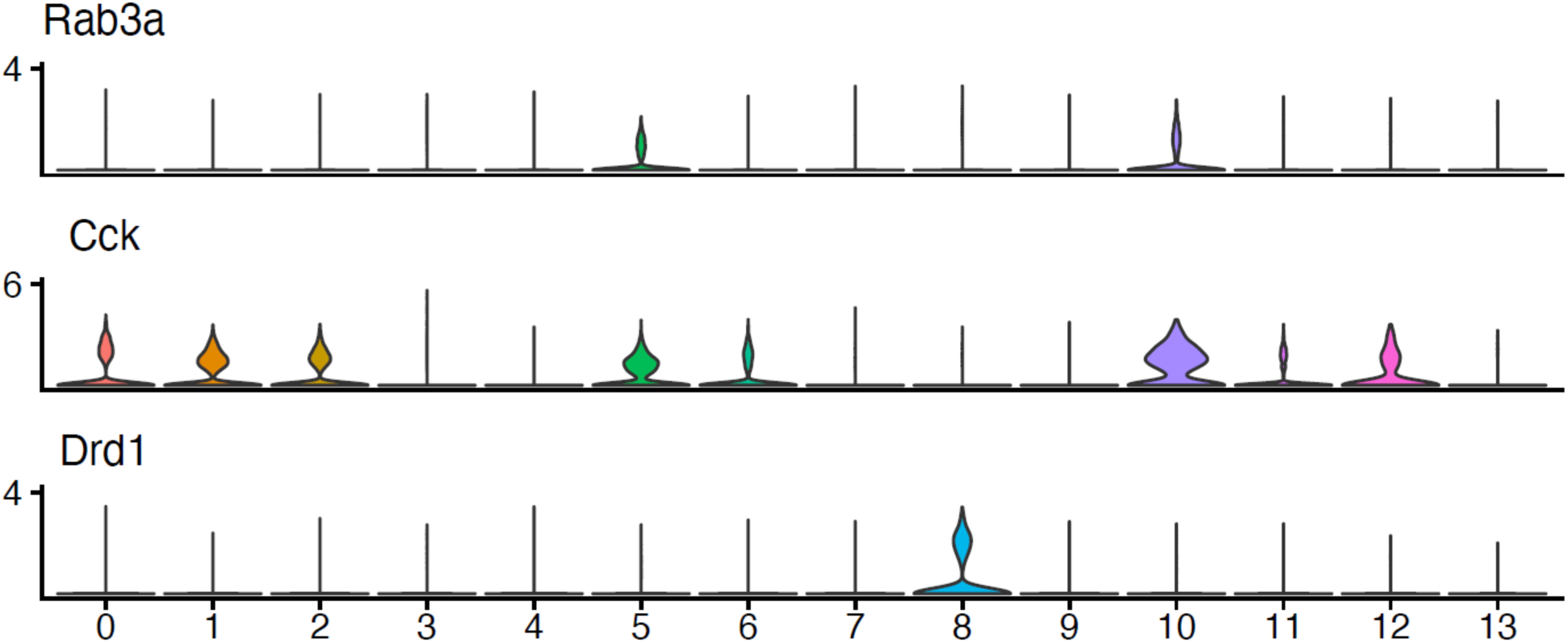
Violin plots showing the level of expression of two mRNAs (*Rab3a* (*24*), *Cck* (*25*)) whose expression levels were shown to be altered during cocaine withdrawal (*19*); and of *Drd1* mRNA, which was primarily expressed in cluster 8. The cluster number is indicated on the x-axis and the y-axis shows the natural log transformed and normalized level of expression for each mRNA.

## Notes

https://www.ncbi.nlm.nih.gov/geo/

### Competing Interest Statement

The authors have declared no competing interest.

